# SF3B1-targeted Splicing Inhibition Triggers Transcriptional Stress Response and Global Alterations in R-Loop Landscape

**DOI:** 10.1101/2020.06.08.130583

**Authors:** Daisy Castillo-Guzman, Stella R. Hartono, Meghan Frederick, Lionel A. Sanz, Tadas Sereiva, Frédéric Chédin

## Abstract

Efficient co-transcriptional splicing is thought to suppress genome-destabilizing R-loops. Inhibition of SF3B1, a core U2 spliceosome component, by Pladienolide B (PladB) in human K562 cells caused widespread intron retention and modest R-loops gains. Minimal overlap existed between these events, suggesting that unspliced introns by themselves do not cause excessive R-loops. R-loop gains were instead driven by extensive readthrough transcription at a subset of stress-response genes, defining a new class of aberrant “downstream of genes” (DoG) R-loops. Such DoG R-loops were temporally and spatially uncoupled from loci experiencing DNA damage. Unexpectedly, the predominant response to splicing inhibition was a global R-loop loss. This resulted from increased promoter-proximal pausing and defective transcription elongation associated with premature termination. Similar results were observed upon depletion of Aquarius, a U2 spliceosome-associated factor previously thought to suppress R-loops. Thus, U2 spliceosome-targeted splicing inhibition triggered profound alterations in transcriptional dynamics, leading to unexpected disruptions in R-loop landscapes.

**HIGHLIGHTS:**

- Intron retention caused by SF3B1 inhibition does not trigger excessive R-loops
- Stress-response genes shows readthrough transcription and R-loop gains
- R-loop gains and DNA damage are temporally and spatially uncoupled
- U2 snRNP inhibition causes broad reduction in transcription and dominant R-loop loss

## INTRODUCTION

During transcription, the nascent RNA can anneal to the DNA template strand behind the advancing RNA polymerase (RNAP), forming a stable RNA:DNA hybrid and causing the non-template DNA strand to loop out. The resulting non-B DNA structure, called an R-loop, is facilitated by favorable DNA sequence and negative superhelicity (Chedin and Benham, 2020; Drolet et al., 2003; Stolz et al., 2019). R-loops have been described from bacteria to plants to mammals (Santos-Pereira and Aguilera, 2015) and represent a prevalent class of alternative DNA structures (Chedin, 2016). Studies over the last decade have implicated R-loop formation in a variety of physiological processes such as transcription termination (Proudfoot, 2016; Sanz et al., 2016; Skourti-Stathaki et al., 2011), chromatin patterning and gene expression control (Chedin, 2016; Niehrs and Luke, 2020; Sanz et al., 2016; Skourti-Stathaki et al., 2019; Tan-Wong et al., 2019), and class switch recombination (Yu et al., 2003; Yu and Lieber, 2019). By contrast, so-called harmful R-loops formed under a variety of pathological conditions have been extensively linked to phenomena of genome instability (Crossley et al., 2019; Garcia-Muse and Aguilera, 2019; Hamperl and Cimprich, 2014). Not surprisingly, a number of human diseases have recently been linked to altered R-loop metabolism (Richard and Manley, 2017). Despite the rising importance of R-loops in adaptive and maladaptive processes, many questions remain regarding the mechanisms regulating their formation and resolution.

It is widely accepted that R-loops form co-transcriptionally upon re-invasion of the nascent RNA *in cis*. Interestingly, alterations of a number of co-transcriptional processes such as splicing, RNA export, cleavage and polyadenylation, have been linked to phenomena of R-loop-mediated genome instability from yeast to human cells (Aguilera, 2005; Aguilera and Garcia-Muse, 2012; Chan et al., 2014; Huertas and Aguilera, 2003; Li and Manley, 2005, 2006; Paulsen et al., 2009; Stirling et al., 2012). Earlier work suggested that inactivation in the splicing regulator SRSF1 triggered the accumulation of R-loops, presumably upon formation of excessive R-loops via newly unspliced transcript portions (Tresini et al., 2015). SRSF1-depleted cells also showed accumulation of double-stranded DNA breaks (DSBs), a hypermutagenic phenotype, G2 arrest, and loss of cell viability (Li and Manley, 2005). Importantly, these phenotypes could be reversed upon over-expression of ribonuclease H1 (RNase H1), an enzyme with the clear biochemical ability to degrade RNA in the context of RNA:DNA hybrids (Cerritelli and Crouch, 2009). This suggested that excessive R-loop formation is directly involved in driving genome instability, and that splicing factors can limit R-loop formation. Systematic genetic screens in human cells identified splicing-related genes as the top category of factors involved in the prevention of R-loop-mediated genome instability, as scored by the phosphorylated H2AX (γH2AX) DNA damage marker (Paulsen et al., 2009). Complementary work in yeast further underscored that introns attenuate R-loop formation and transcription-associated genetic instability via the recruitment of the spliceosome onto the pre-mRNA (Bonnet et al., 2017). Altogether, these studies suggest that proper splicing, export, and packaging of the nascent RNA into ribonucleoprotein particles contributes significantly to the maintenance of genome stability by keeping the RNA away from the DNA template and avoiding deleterious R-loop formation.

The nature and distribution of excessive R-loops expected to result from the disruption of co-transcriptional RNA processing have not been established at genome scale. Thus, the spatial relationship between putative aberrant R-loop gains and events of genome instability has never been directly tested. To address these gaps, we focused on splicing as a key co-transcriptional process linked to R-loop regulation. SF3B1 is a conserved and essential core subunit of the SF3B complex, a key component of the U2 spliceosome involved in branch site recognition and selection during pre-mRNA splicing (Sun, 2020). SF3B1 can be specifically inhibited via pharmacological compounds, including Pladienolide B (PladB) (Cretu et al., 2018; Kotake et al., 2007; Yokoi et al., 2011). PladB and other macrolide-type splicing inhibitors rapidly abrogate splicing, causing a vast increase in events of intron retention (IR) (Boswell et al., 2017; Carvalho et al., 2017; Kashyap et al., 2015). PladB treatment provides a unique opportunity to determine whether unspliced introns can drive excessive R-loop formation and associated DNA damage, which is expected if the newly retained intron sequences form genome-destabilizing R-loops. This pharmacological approach further allowed us to temporally resolve the resulting changes in R-loop distribution, splicing events, and transcriptional dynamics, providing a unique view into the cellular response to acute splicing inhibition.

## RESULTS

### PladB treatment causes broad intron retention and R-loop reduction in human K562 cells

We used RNA-seq to characterize alterations in splicing patterns resulting from PladB treatment two and four hours after drug application. Human K562 cells either mock-treated with DMSO or untreated were used as controls and two biological replicates were analyzed at each time point (Figure 1A). As expected, PladB treatment caused broad, time-dependent, splicing alterations dominated by events of intron retention (IR) and to a lesser extent, skipped exons (SE) (Figure 1B). Validation of RNA-seq analysis by RT-qPCR at the known PladB target *DNAJB1* (Kashyap et al., 2015) confirmed that IR gradually accumulated over time and could be detected as soon as 15 minutes after treatment (Figure S1A,B). IR was observed at *DNAJB1* in strand-specific RNA-seq analysis of poly(A)-tailed mRNA pools two- and four-hours post treatment (Figure 1C). In total, over 7,000 independent IR events were identified in human K562 cells four hours post PladB treatment. Such retained introns provide an ideal cohort to test the hypothesis that splicing inhibition causes gains of R-loops over regions now associated with unspliced introns. To determine how R-loops respond to PladB-mediated splicing inhibition, we profiled R-loop distribution using DRIP-seq (Sanz and Chedin, 2019) at the same time points. Only a modest number of R-loop gains (RLGs) were observed post PladB treatment with 1,467 and 1,878 peaks identified within the limits of our statistical thresholds two and four hours post-PladB, respectively (Figure 1D). Unexpectedly, the vast majority of significant R-loop changes corresponded to events of R-loop loss, highlighting 18,150 and 28,844 peaks two and four hours post-PladB, respectively. Most loci did not show significant R-loop gains or losses.

**FIGURE 1:**
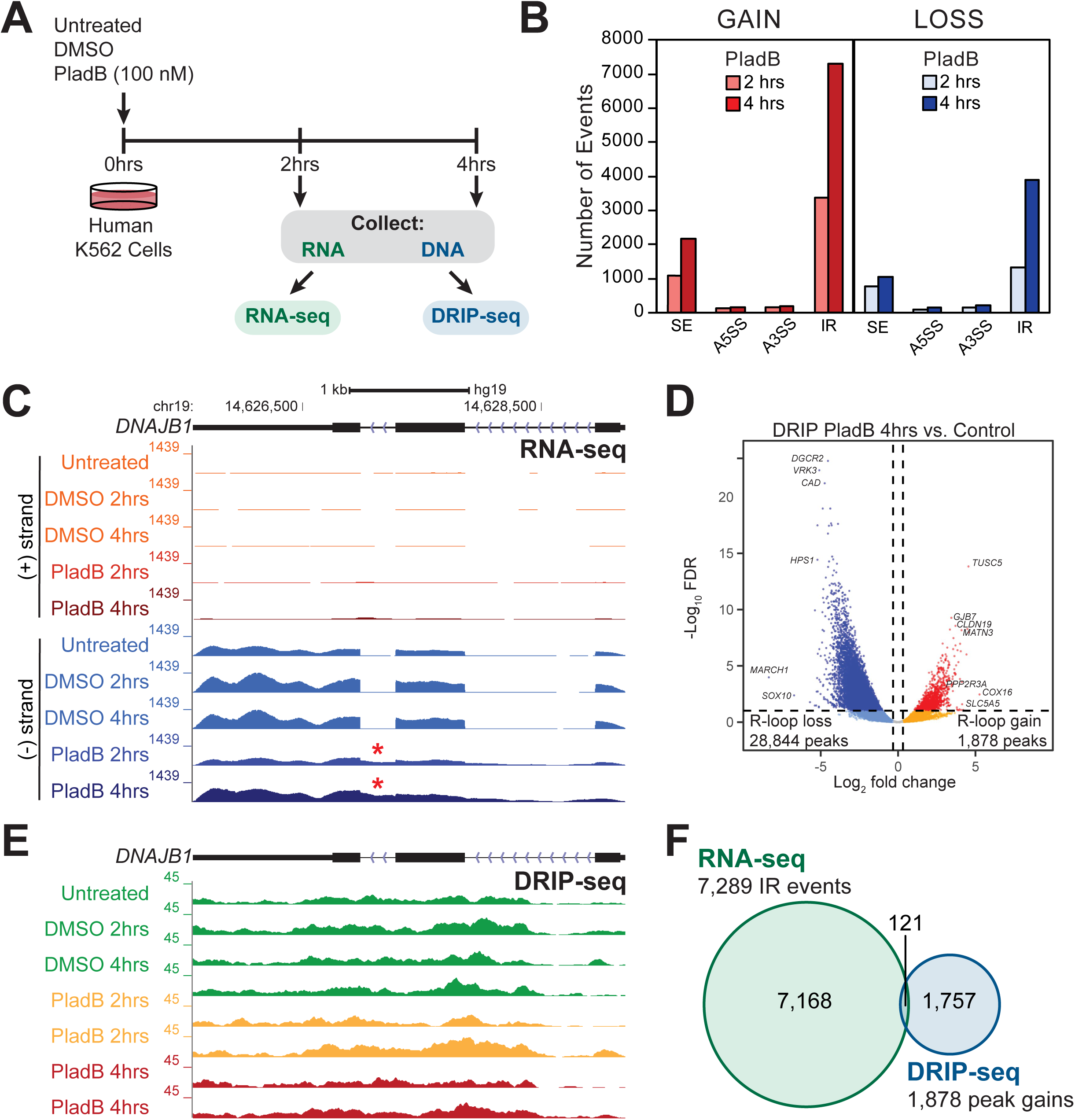
**A.** Schematic of experimental setup, see text for details. **B.** Summary of PladB-mediated splicing alterations identified by RNA-seq. The number of skipped exons (SE), alternative 5’ splice sites (A5SS), alternative 3’ splice sites (A3SS) and intron retention (IR) gained or lost after 2 and 4 hours of PladB treatment are graphed. **C.** Genome browser screenshot over the representative *DNAJB1* gene showing plus and minus strand poly(A) RNA-seq signal obtained from controls and PladB treated K562 cells. Intron retention is observed two- and four-hours post PladB treatment (asterisk). **D.** XY plot of normalized DRIP-seq counts for control and treated (PladB 4 hours) samples. Each dot on the graph corresponds to a DRIP-seq peak. Red/blue colors indicate significant gains/losses of signals (adjusted p-value of < 0.05 and |log2(FC)| ≥ 1). **E.** Genome browser screenshot showing the same region as in (C) but now displaying DRIP-seq signals. No increase in R-loop signal is detected over the retained intron. **F.** Venn diagrams displaying the intersection of events of intron retention and R-loop gains.

To determine whether IR associates with RLGs, we first focused on *DNAJB1* that showed clear intron retention: no significant RLGs could be observed (Figure 1E). R-loops were in fact significantly reduced at the *RPL13A* gene despite clear IR (Figures S1C, S1D). More broadly, less than 2% of loci with IR events intersected with RLG peaks (121 **/** 7289; Figure 1F). Thus, contrary to expectations, large-scale intron retention caused by acute PladB-mediated splicing inhibition, did not cause R-loop gains over unspliced transcripts.

### Splicing inhibition is associated with readthrough transcription and accompanying R-loops at a subset of genes

To understand the nature of RLGs caused by PladB, we analyzed the distribution of these peaks in the genome. 77% of R-loop peaks in untreated K562 cells mapped to genic regions, as expected (Sanz et al., 2016). By contrast, nearly two thirds of RLGs in PladB-treated cells mapped to intergenic regions (Figure 2A), indicating that PladB treatment unexpectedly led to RLGs over non-genic regions. To assess how such gains arose, we focused on a region with numerous clustered RLGs peaks located downstream of the *VAPA* gene (Figure 2B). The peaks extended from the 3’-end of *VAPA* and grew directionally with transcription in a time-dependent manner. After two hours, the edge of the wave of RLGs extended 135 kilobases (kb) downstream of the *VAPA* gene. After four hours, a trail of R-loops was detected up to 210 kb downstream of *VAPA*. DRIP-qPCR on three additional replicates confirmed that R-loops increased 20-fold 60 kb downstream of the *VAPA* poly(A) site (PAS) and 6-fold at a region located 200 kb downstream of *VAPA* four hours post treatment. All instances of R-loops were sensitive to RNase H pre-treatment, as expected (Figure 2C). This pattern is consistent with a directional and time-dependent propagation of a wave of co-transcriptional R-loops from the *VAPA* gene. Analysis of R-loops using a high-resolution, strand-specific, iteration of DRIP-seq (sDRIP-seq; see methods) confirmed that the RLGs were stranded and co-directional with *VAPA* transcription (Figure S2A). Similar findings were observed for two additional representative genes, *RPL9* and *CYCS* that showed a stranded increase in R-loop signal extending downstream of their PAS (Figure S2B-D). Given that R-loops form co-transcriptionally, these observations suggest that PladB-mediated splicing inhibition caused readthrough transcription downstream of *VAPA*. RT-qPCR analysis of total RNA confirmed the existence of transcripts downstream of *VAPA* upon PladB treatment (Figure 2D). Transcripts were detected 15 and 60 kb downstream of *VAPA* 30 min post PladB treatment and steadily built over time up to a 35 to 60-fold increase over controls. Far downstream of *VAPA* (120 kb), transcripts were only detected 120 and 240 min after treatment, consistent with the steady progression of the RNA polymerase (RNAP) from the 3’- end of *VAPA* and not from spurious transcription initiation in intergenic regions. To further explore the possibility that PladB treatment triggers downstream of genes (DoG) transcription, we analyzed nascent transcripts in control and PladB-treated K562 cells using EU labeling and RNA sequencing of labeled transcripts. Nascent transcripts were clearly observed downstream of *VAPA* and progressed unidirectionally as a function of time from the normal *VAPA* termination region (Figure 2E). Similar findings were observed at the *CYCS* and *RPL9* genes (Figure S2E-F). Thus, PladB treatment triggered readthrough transcription and accompanying co-transcriptional R-loops.

**FIGURE 2:**
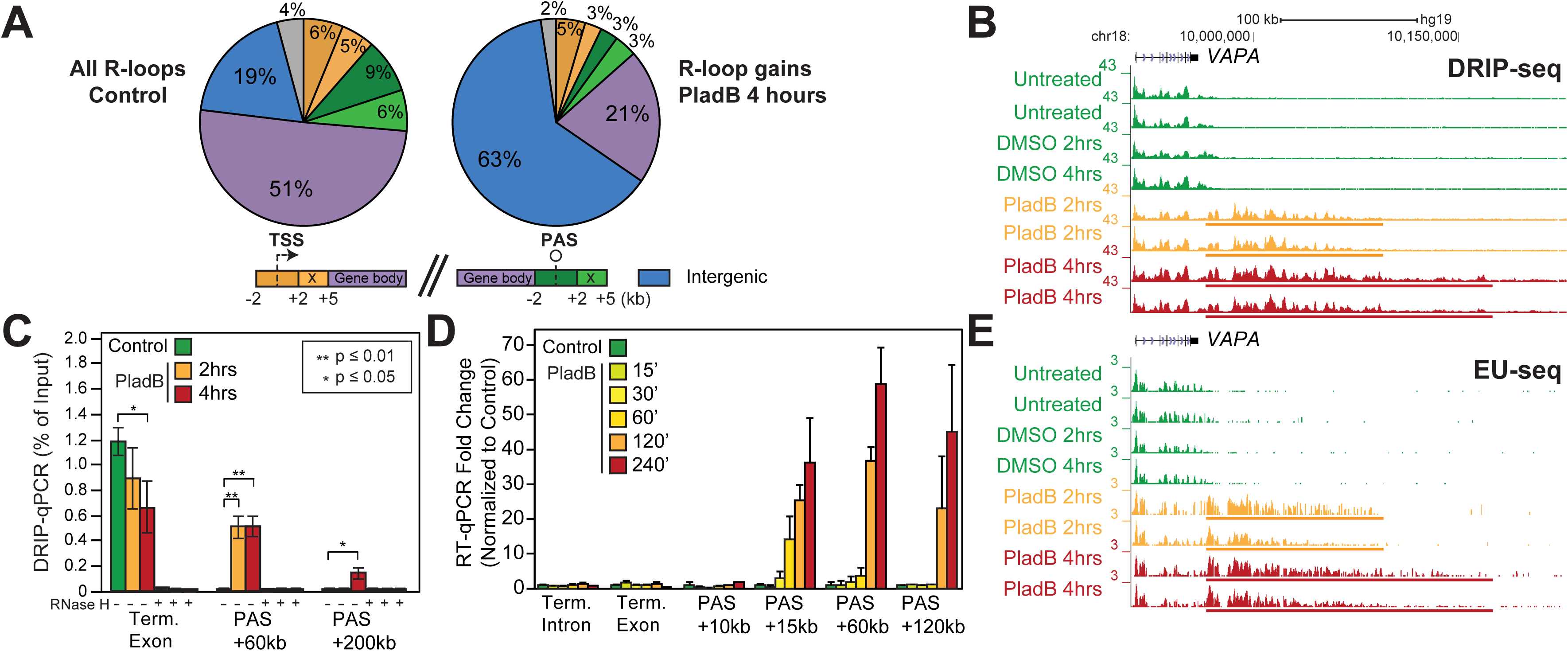
**A.** Location analysis of DRIP peak gains for control (left) and PladB-treated samples (right) across genomic compartments (color-coded according to the schematic below). R-loop distribution across compartments is indicated by percentages. TSS, transcription start site; PAS, polyadenylation site. **B.** Genome browser screenshot over the representative *VAPA* gene and ∼200 kb downstream showing DRIP-seq signal from controls and PladB-treated K562 samples. R-loop gains occur directly downstream of gene (DoG - colored bar). **C.** Bar chart of DRIP-qPCR (as percent of input) for *VAPA* and two regions downstream of PAS. Each bar is the average of three-independent experiments (shown with SE). **D.** Bar chart of RT-qPCR (as fold change over control) at indicated loci through a time course after PladB treatment. Each bar represents the average of three independent experiments (shown with SD). **E.** Genome browser screenshot over the same region as in (B) but now displaying EU-seq signals. Regions with increased R-loops also have increased transcription.

### DoG transcription affects a specific subset of stress-responsive genes and is associated with broad *de novo* R-loop gains

To determine how broad the PladB-induced transcriptional readthrough was, we systematically annotated whether events of RLGs correspond to DoG R-loops initiating from a neighboring upstream gene. A total of 429 genes, referred to thereafter as DoG genes, were identified with these characteristics, corresponding to the large majority of RLGs (1,505/1,878 peaks; Figure S3A). Metaplots confirmed increased R-loops and nascent transcripts extending downstream of the host genes PASs for a distance of 50-75 kb (Figure 3A, B; Figure S3B). Annotation of individual DoG R-loops confirmed that the median length of DoG R-loop regions was 49.2 and 50 kb two- and four-hours post PladB treatment, respectively (Figure 3C). This was significantly higher than the length of terminal R-loops recorded for the same genes under control conditions. Interestingly, PladB-sensitive DoG genes showed longer terminal R-loop regions than expression-matched genes even under control conditions (31.5 kb versus 18.2 kb) indicating that DoG genes may have an intrinsic propensity to terminate further downstream of their PASs (Figure 3C). Collectively, DoG R-loops observed four hours after PladB treatment covered 2.81 megabases of genomic space, defining a class of excessive R-loops at genome-scale. Interestingly, the propensity for R-loop formation, measured by plotting the ratio of DRIP signal over nascent transcript signal, was significantly higher in DoG regions compared to gene bodies (Figure S3D) even though there was no measurable increase in R-loop-favorable characteristics such as GC skew or GC content over DoG regions. Thus, DoG transcription appears to be prone to R-loop formation.

**FIGURE 3:**
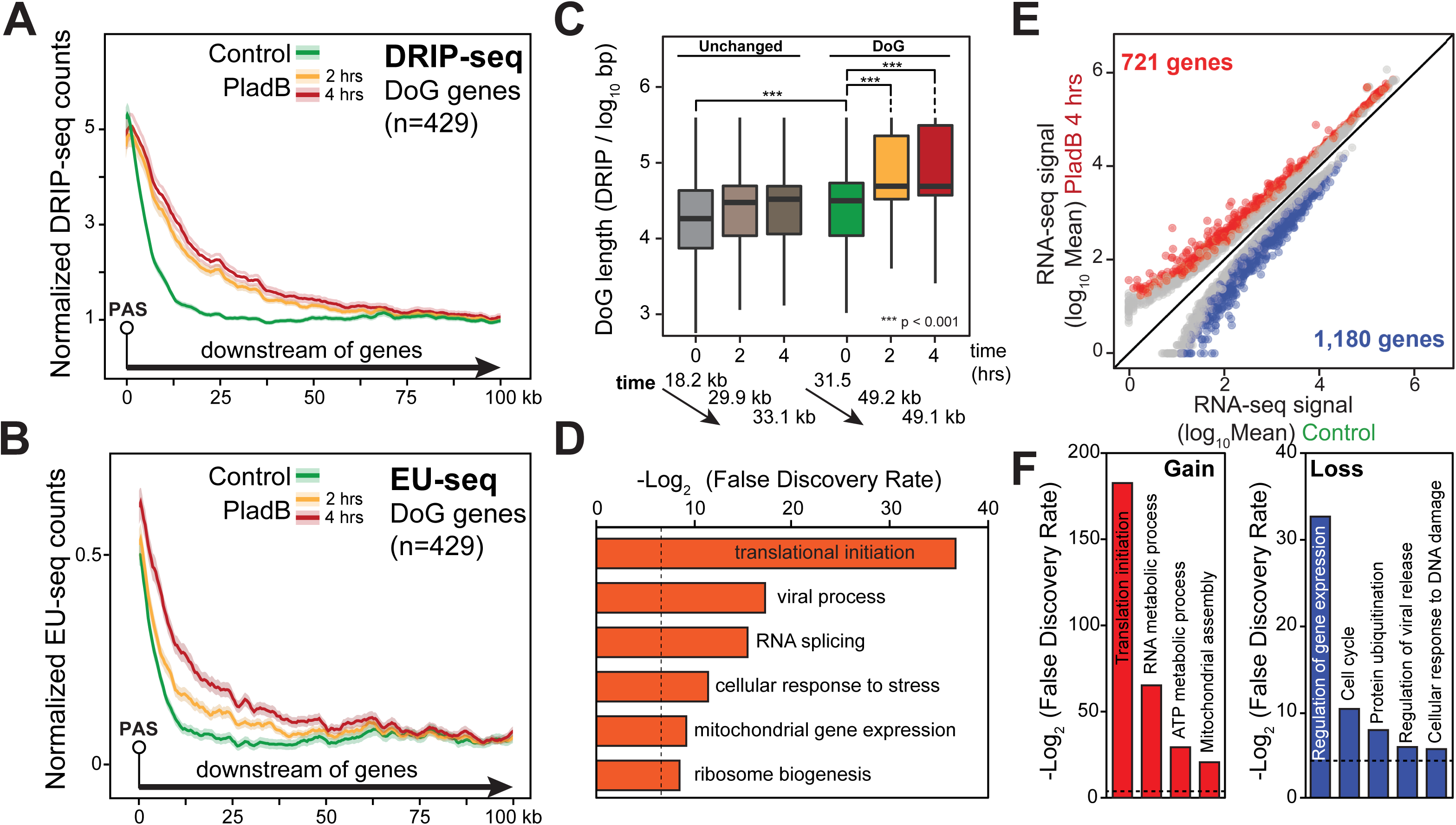
**A,B.** Metaplots of DRIP-seq (top) and EU-seq (bottom) signals extending from nearest PAS for all annotated DoG genes. For each time point, the signal is shown as a trimmed mean (line) surrounded by SE (shaded). **C.** DoG R-loop lengths (measured from DRIP-seq data) under control conditions and two- and four-hours post PladB for DoG genes and expression-matched genes that showed no significant R-loop changes. P-values were determined by a Wilcoxon Mann-Whitney test. The numbers below indicate median terminal R-loop lengths at each time points. **D.** Enriched gene ontologies for DoG genes. The dashed line indicates 1 % FDR. **E.** XY plot of normalized RNA-seq counts for control and treated (4 hours PladB) samples. Each dot on the graph corresponds to a gene. Red/blue colors indicate gains/losses of RNA-seq signals (adjusted p-value of ≤ 0.05 and |log2(FC)| ≥ 1). **F.** Enriched gene ontologies for RNA-seq gain genes and for RNA-seq loss genes. The dashed line indicates 5% FDR.

DoG genes showed significant enrichments for specific gene ontologies (GOs), including translation initiation and ribosome biogenesis, viral process, response to stress, mitochondrial gene expression, and chromatin organization (Figure 3D, Figure S3C). The appearance of stress-related and viral-related GOs is notable given precedents for DoG transcription triggered upon environmental stresses and viral infections (Bauer et al., 2018; Cardiello et al., 2018; Hennig et al., 2018; Vilborg et al., 2015; Vilborg et al., 2017). To investigate if DoG genes arise in part due to elevated transcription, we measured changes to mRNA pools following PladB treatment using RNA-seq. 721 genes showed significant increases in gene expression (Figure 3E). These up-regulated genes were dramatically enriched for a few, highly congruent, biological functions, including protein translation and ribosome biogenesis, RNA metabolic processes including RNA splicing, and mitochondrial ATP synthesis (Figure 3F). Similar ontologies were observed for DoG genes (Figure 3D), suggesting that R-loop gains reflected at least in part an increase in gene expression. Another 1,180 genes were significantly down-regulated and displayed enrichment for functions related to gene expression regulation, cell cycle control, protein ubiquitination, regulation of viral release, and the cellular response to DNA damage (Figure 3F). Overall, PladB triggered a strong cellular stress response associated with DoG transcription and accompanying DoG R-loops for a subset of stress-responsive genes.

### Global R-loop losses are caused by PladB-induced negative feedback on transcription elongation

The overwhelming majority of R-loop changes in response to PladB (28,844 / 30,713 events, or 93.8%, 4 hours post PladB) unexpectedly corresponded to R-loop losses (Figure 1D). To understand the mechanisms driving these events, we first asked where R-loop losses (RLLs) occurred. Contrary to RLGs, RLLs matched primarily to transcribed gene bodies (44%) and terminal genic regions (25%) (Figure 4A). An additional 20% of RLLs matched to normally transcribed regions located immediately downstream of terminal PAS sites. By contrast, only a small minority of RLLs mapped to promoter and promoter-proximal regions. To further understand how these losses originated, we focused on the representative R-loop-prone *DGCR2* gene (Figure 4B). RLLs were clearly visible through the gene body, with little initial impact around the promoter region and early first intron. These RLLs were validated by DRIP-qPCR in three additional independent replicates, which confirmed that early R-loops were unchanged but R-loop formation along the gene body was progressively reduced from 10-fold in the middle of the gene to 50-fold by the gene terminus (Figure 4C). The simplest hypothesis to account for these observations is that transcription elongation was impaired, leading to a directional reduction of co-transcriptional R-loops. To test this, we took advantage of our matched EU-seq datasets and observed that nascent transcripts along *DGCR2* were significantly affected, with EU incorporation only located in the early portion of intron 1, where R-loops still occurred (Figure 4D). Similar observations were made for other genes and further validated by DRIP-qPCR and sDRIP-seq (Figure S4A-D).

**FIGURE 4:**
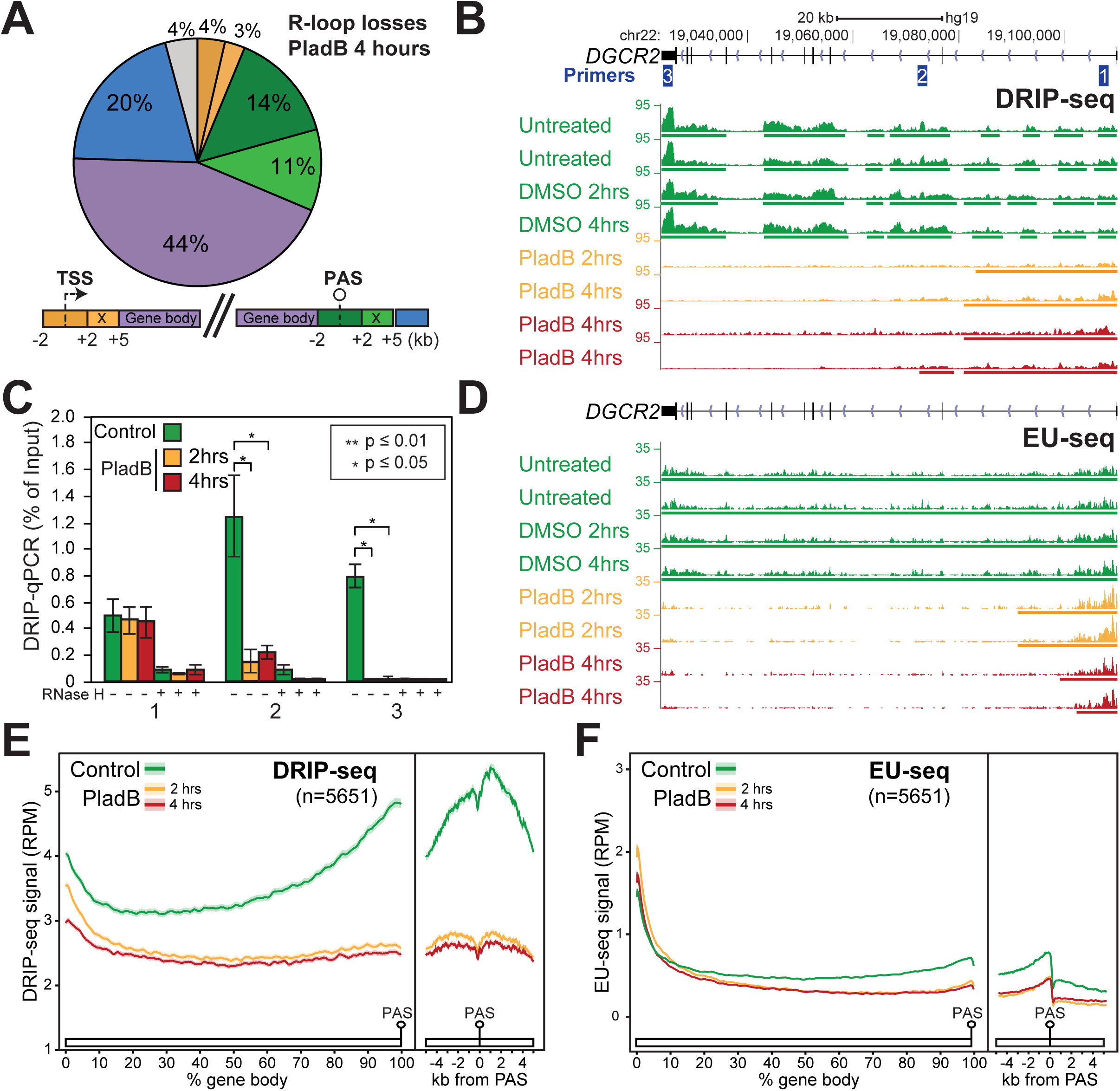
**A.** Location analysis of R-loop losses for PladB-treated (4 hours) samples across genomic compartments. Numbers indicate the percentage occupied by each compartment. **B.** Genome browser screenshot over the representative *DGCR2* gene showing DRIP-seq signal from controls and PladB-treated K562 samples (colored bars indicate R-loop peaks). **C.** Bar chart of DRIP-qPCR (as percent of input) for *DGCR2* using indicated qPCR primer pairs. Each bar is the average of three-independent experiments (shown with SE). **D.** Genome browser screenshot showing the same region as in (B) but now displaying EU-seq signals. **E.** Metaplot of DRIP-seq signals over gene body (as % of gene length) and terminal regions (+/- 5kb from PAS) of EL genes. Control and PladB-treated samples are color-coded as indicated. For each time point, the signal is shown as a trimmed mean (line) surrounded by the standard error (shaded). **F.** Same as E except EU-seq signals are plotted.

We systematically annotated genes showing significant RLL in their gene body or terminal regions and identified 5,651 genes, referred to thereafter as RLL genes. As expected from single gene examples, metaplot analyses confirmed loss of R-loops gradually occurred along gene bodies and was maximal around gene termini (Figure 4E; Figure S4E). Clear loss of transcription was also observed along these genes (Figure 4F). In general, nascent transcripts and R-loops were in strong agreement: 77% of genes independently annotated as showing loss of nascent transcripts also showed RLL. The remainder of genes showing EU-seq loss without accompanying RLL simply had little to no R-loop signal under control conditions. Conversely, about half of the RLL genes also showed loss of transcription. Overall, this data suggests that splicing inhibition triggered profound negative feedback on transcription elongation, resulting in a global reduction of co-transcriptional R-loops.

Our findings thus far indicate that PladB treatment resulted in contrasting alterations in transcriptional dynamics for different gene sets. A large subset of RLL genes showed transcription elongation and R-loop losses in response to PladB. By contrast, a smaller group of DoG genes showed readthrough transcription and associated R-loop gains. In addition, we identified a third large class of transcribed genes (n=5,455) that showed no significant loss of gene body R-loops, nor gains of DoG R-loops. These genes will be referred to as “unchanged” and represent a useful internal control. We next sought to identify the mechanisms leading to transcription elongation loss and asked if the gene classes identified above possess differential characteristics that could account for their unique responses to PladB.

### Splicing inhibition triggers global premature termination

Inhibition of U1 small nuclear ribonucleoprotein (snRNP) triggers strong negative feedback on transcription caused by premature termination via the cleavage and polyadenylation machinery (Kaida et al., 2010). We investigated if a similar effect was observed upon PladB treatment by annotating putative PladB-specific alternative polyadenylation (APA) sites in RNA-seq data. The *BRD4* gene, which regulates the release of promoter-paused RNA polymerase complexes as well as transcription elongation and termination (Aoi and Shilatifard, 2023; Arnold et al., 2021; Winter et al., 2017), showed strong APA up-regulation (Figure 5A). In mock-treated cells, transcription led to the expression of two long *BRD4* isoforms, as expected (Han et al., 2020). PladB led to significant reduction in the longest isoform and the up-regulation of a short annotated isoform ending after exon 11. In addition, a novel 3’-UTR ending at an annotated PAS site after exon 7 was sharply induced by PladB. RT-qPCR experiments confirmed that the abundance of RNAs overlapping this novel putative APA increased 15-fold 4 hours post PladB treatment, while the long *BRD4* isoform was reduced 2-fold (Figure S5A). Both exon 7 and 11 isoforms ended at or immediately adjacent to an annotated polyadenylation site (PAS) (Figure 5A). Thus, BRD4, a critical transcriptional regulator undergoes striking PladB-induced APA, possibly leading to the expression of short BRD4 isoforms lacking much of the BRD4 C-terminal domain.

**FIGURE 5:**
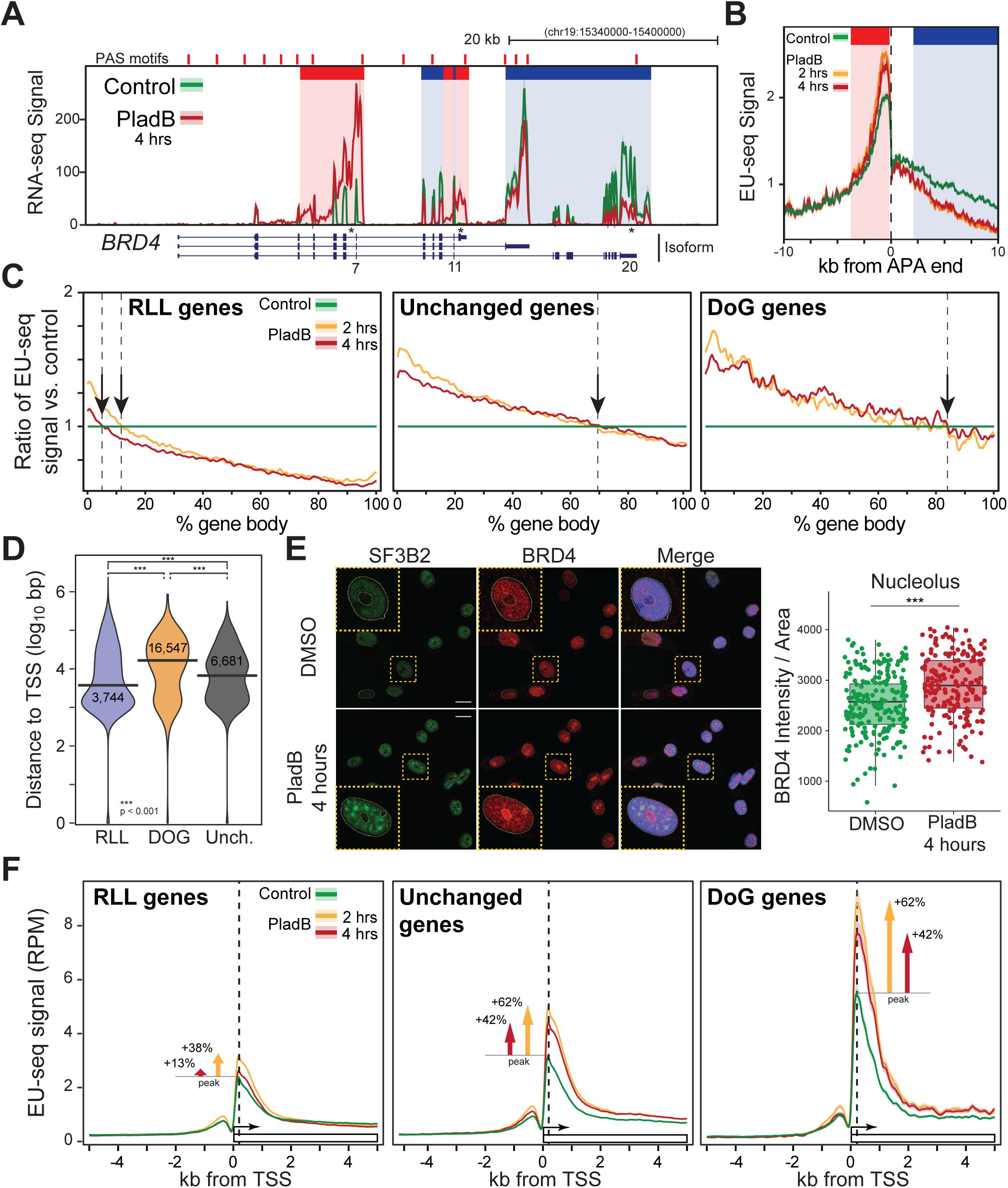
**A.** RNA-seq signal for *BRD4* under control and PladB-treated conditions. Highlighted in red and blue are sites of significantly increased or decreased APA usage (adj. p-value < 0.05). **B.** Metaplot of EU-seq signals under control and PladB-treated conditions. Reads were aligned at the end of novel, PladB-specific, intronic 3’-UTRs (+/- from APA end). **C.** Ratio of mean EU-seq signals (PladB/control) for DoG, unaffected, and RLL genes plotted over gene bodies for control and PladB-treated conditions. The positions at which PladB nascent transcript signals fall below control signals are highlighted by vertical arrows. **D.** Distance from the TSS to the first PladB-induced intronic 3’-UTR for DoG, unaffected, and RLL genes. **E.** Representative immunofluorescence microscopy images from DMSO-and PladB-treated (4 hours) HeLa cells analyzed for SF3B2 and BRD4 distribution and DAPI (scale bar 10 μm). Quantification of the BRD4 immunofluorescence signal over nucleoli; each dot represents an individual cell measured across 2 biological replicates per condition. **F.** Metaplot of EU-seq signals over DoG, unaffected, and RLL gene promoters (+/- 5 kb from TSS) under control and PladB-treated conditions. The relative signal increases measured at the promoter-proximal peak and 2 kb downstream of TSS are indicated.

Systematic annotations revealed 5,771 putative APA sites 4 hours after PladB treatment. Half of these events mapped onto previously annotated 3’UTRs, while the other half defined novel putative intronic APAs (Figure S5B). These candidate intronic APAs were in close proximity to poly(A) motifs (Figure S5C), suggesting they correspond to *bona fide* APA sites. Representative examples from the dataset show clear PladB-specific APAs that were validated by RT-qPCR (Figure S5A, D). Overall, PladB-induced APA events mapped onto 3,631 unique genes showing gene ontology enrichment similar to those observed for down-regulated genes including RNA splicing, cell cycle regulation, the response to DNA damage, and the IRE1-induced unfolded protein response (Figure S5E). Overall, PladB-mediated U2 spliceosome inhibition led to premature termination and profound changes in APA patterns, causing widespread isoform switching.

Increased premature termination should lead to reduced nascent transcripts downstream of APA sites. In agreement, 80% of the genes showing premature APA usage (2,869 / 3,631 genes) showed significant losses in transcription (EU-seq), gene expression (RNA-seq), and/or R-loop levels (DRIP-seq). Metagene plots of EU-seq signal centered at the ends of PladB-specific intronic APAs confirmed increased stalling over the APAs followed by decreased transcription immediately downstream, consistent with termination (Figure 5B).

To determine if the three gene classes responded differentially to PladB-induced APA, we plotted the ratios of EU-seq signals observed two and four hours after PladB treatment over the EU-seq signal for the same genes under control conditions (Figure 5C). RLL genes showed a minor increase in nascent transcripts very early in gene bodies, followed by a steady decline such that the transcription output under PladB treatment fell below that of control cells within the first ∼10% of gene bodies. In contrast, unchanged genes showed a strong increase in nascent transcripts immediately downstream of the TSS such that the transcription output under PladB only fell below that of control cells after about 70% through gene bodies. DoG genes had a similar behavior, except the stranscription output under PladB fell below control only towards the end of these genes. PladB therefore caused a gradual elongation defect in all three gene classes, as visualized by similar negative slopes in EU-seq ratios. However, the large increase in transcription observed early in gene bodies for unchanged and DoG genes partially compensated for decreased elongation and allowed a significant number of RNAPII complexes to reach the end of genes. RLL genes, by contrast, suffered dramatic transcription loss early in elongation. To further address if the reduced elongation was linked to premature APA, we annotated the distance from the TSS to PladB-induced intronic APA events in each gene class.

APA events were located in close proximity downstream of the TSS for RLL genes (median distance 3.7 kb, Figure 5D). In sharp contrast, DoG genes showed a >4-fold greater median distance between their TSS and intronic APAs (median distance 16.5 kb). Unaffected genes showed intermediate distances. Thus, while all gene classes suffered from PladB-induced premature transcription termination, RLL genes appear particularly vulnerable to it because APA events were encountered significantly earlier during elongation.

### PladB treatment leads to increased promoter-proximal pausing

The observation that *BRD4* underwent premature APA suggested that BRD4 protein levels could be reduced by PladB. Western blots and mass spectrometry assays, however, did not reveal any significant change in overall BRD4 levels over short time intervals (2 and 4 hours). Imaging assays instead revealed that the sub-cellular distribution of BRD4 was significantly altered upon splicing inhibition, with clear BRD4 redistribution towards nucleoli (Figure 5E). SF3B2, as expected, was retained in enlarged splicing speckles. Nucleolar sequestration may therefore reduce functional BRD4 pools and affect its ability to mediate pause release at RNAPII promoters. In addition, cyclin T1 also became localized to nuclear speckles upon PladB treatment (Figure S5F), possibly reducing functional levels of the critical pause release factor, pTEF-b. We therefore examined patterns of nascent transcripts in the vicinity of TSSs as a proxy for pausing. DoG genes showed the most pronounced EU-seq signal immediately downstream of the transcription start site (TSS) (Figure 5F, right). This peak was sharply reduced over the first 2,000 bp of promoter-downstream sequences, suggesting that DoG genes are heavily regulated by pause-release mechanisms and carry high loads of slow moving and/or rapidly recycled RNAPs early in their gene bodies. Traveling ratios (TRs) (Rahl et al., 2010) confirmed that DoG genes showed significantly higher TRs compared to both unchanged and RLL genes under control conditions (Figure S5G). Importantly, PladB treatment caused a further significant increase in DoG gene promoter-proximal pausing. In sharp contrast, RLL genes had the lowest pausing under control conditions and showed the least gains upon PladB treatment (Figure 5G, left). Unchanged genes were more similar to DoG genes in that they showed a strong increase in pausing upon PladB treatment. Thus, the three gene classes defined here solely based on their R-loop behaviors showed important differences in their amounts of promoter-proximal paused RNAP complexes and in their responses to splicing inhibition. Overall, the data suggests that RLL genes are uniquely vulnerable to transcription elongation loss due to their sensitivity to premature APA and their lack of a robust promoter store of paused RNA pol II complexes. In addition, we propose that U2 spliceosome inhibition exerts a negative feedback on RNA polymerase pause release mechanisms, via the sequestration of BRD4 and cyclin T1 in distinct sub-nuclear compartments.

### DoG genes preferentially show distal splicing alterations which may be coupled to readthrough transcription

At first glance, DoG and unchanged genes responded similarly to PladB; yet readthrough transcription and DoG R-loops were only seen for DoG genes. Since proper transcription termination has been linked to terminal exon splicing (Davidson and West, 2013; Herzel et al., 2017; Rigo and Martinson, 2008, 2009), we asked whether DoG genes showed a distinct pattern of splicing alterations. DoG genes showed a pronounced shift of skipped exon events towards the end of gene bodies compared to both RLL and unchanged genes (Figure S5H). Such events often affected terminal or near terminal exons. Similar trends were observed for intron retention events. Interestingly, no single exon gene was present in the DoG gene dataset even though 1,247 single exon genes were expressed in K562 cells and 40 were expected to display DoG transcription at random. These results suggest that DoG genes are significantly more predisposed to aberrant terminal splicing events.

We also wondered if DoG genes might display characteristics predisposing them to weaker transcription termination efficiencies compared to unchanged genes. Analysis of PAS motifs did not reveal significant differences between gene classes. However, DoG genes showed unusual R-loop patterns through their gene bodies under control conditions. As expected from their high loads of promoter-proximal RNAPs, DoG genes showed significantly higher R-loop levels over promoter regions and early gene bodies (Figure S5I). However, DoG genes showed significantly reduced R-loop levels towards gene ends compared to other genes. Unchanged genes, by contrast, showed a clear rise in terminal R-loops, which has been associated with efficient transcription termination (Proudfoot, 2016; Sanz et al., 2016). Overall, the tendency of DoG genes to experience distal splicing alterations upon PladB treatment combined with intrinsically poorer termination mechanisms may predispose DoG genes to readthrough transcription.

### PladB treatment triggers mild DNA damage induction that is temporally and spatially uncoupled from DoG R-loop accumulation

PladB treatment triggers a DNA damage response which has been linked to excessive R-loop formation (Nguyen et al., 2019). We showed here that elevated R-loop levels are detected over DoG genes two hours post-PladB treatment and that DoG transcripts can be detected as early as 30 minutes (Figures 2, 3). We therefore determined if DNA damage induction, as measured by γH2AX immunofluorescence staining, was temporally coupled to DoG R-loop formation. No significant γH2AX foci increase was observed four hours post PladB despite abundant evidence for DoG R-loops. γH2AX nuclear intensity and foci numbers were significantly higher 12 hours post-PladB treatment and increased until 24 hours (Figure 6 A,B; Figure S6A); however, it remained mild compared to the strong positive control, Etoposide (ETO). DNA damage induction was also clearly observed at 24 hours upon staining with the DNA double-strand break marker 53BP1 (Figure S6B). A characteristic gain in nuclear area was also observed over time and by 24 hours cells displayed a well-documented G2/M arrest (Figure S6C,D). Overall, this suggests that the DNA damage triggered by PladB is temporally uncoupled from DoG R-loop accumulation. This delay could be explained if DoG R-loops only occur outside of S phase such that cells do not experience the transcription-replication conflicts thought to drive R-loop-mediated genome instability (Garcia-Muse and Aguilera, 2019). To address this, cells were synchronized (Figure S6E) and treated with PladB while in G1, S, and G2. R-loop levels were then assessed at representative loci for both DoG and RLL genes using DRIP-qPCR. This revealed that PladB-mediated DoG R-loops increases and R-loop losses at RLL genes clearly occurred at all phases of the cell cycle, including S phase (Figure 6C). Thus, the lack of γH2AX foci 4 hours post PladB treatment is likely not due to the absence of DoG R-loops in S phase.

**Figure 6:**
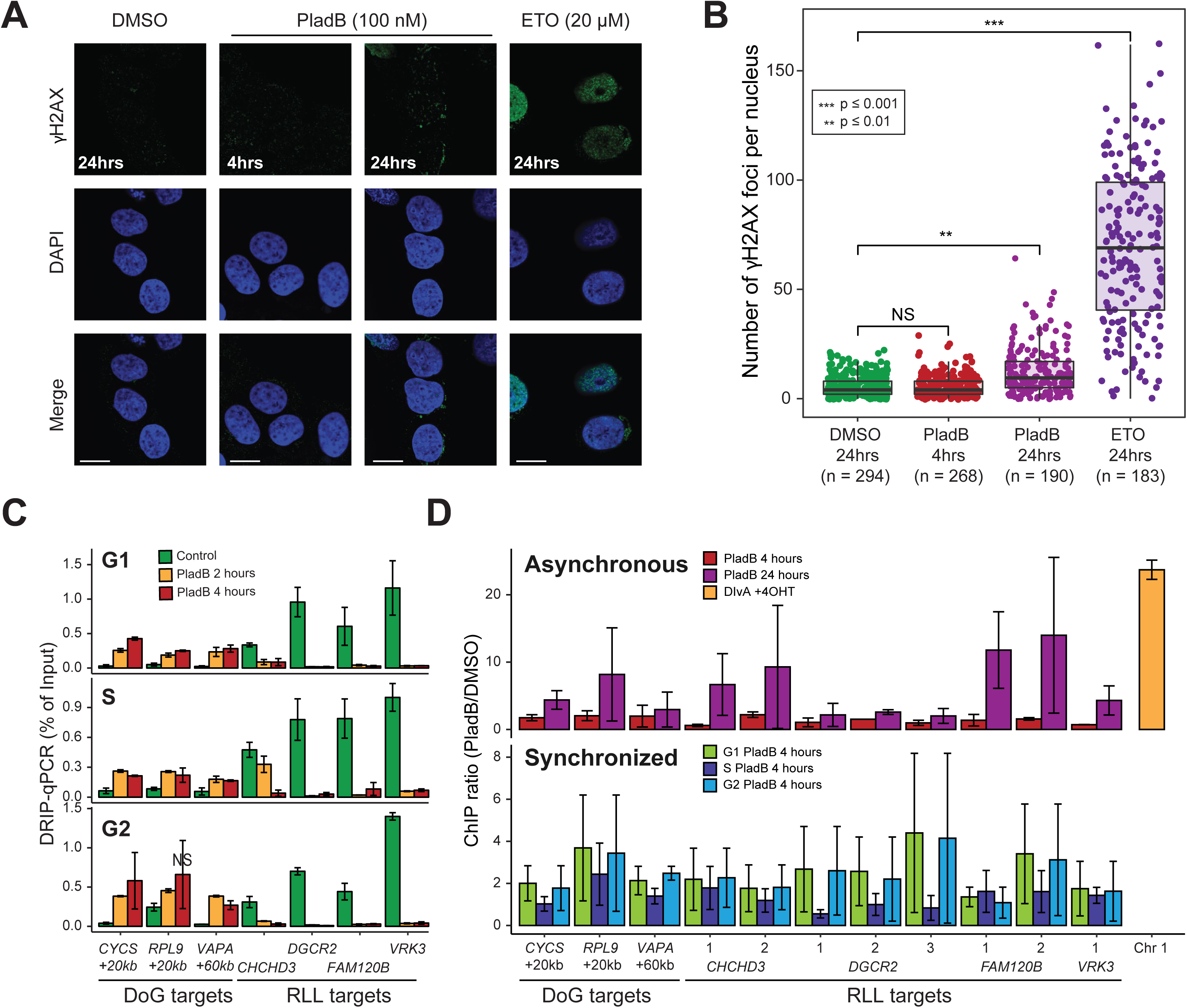
**A.** Representative immunofluorescence microscopy images from DMSO- and PladB-treated (4 and 24 hours), and Etoposide-treated (ETO) HeLa cells analyzed for γH2AX distribution and DAPI. Scale bar 10 microns. **B**. Quantification of the immunofluorescence data for each condition; the number of cells across 2 biological replicates analyzed per condition is indicated. **C**. Bar chart of DRIP-qPCR (as percent of input) for DoG genes and RLL genes measured in cells synchronized in G1, S, or G2/M. Each bar is the average of two-independent experiments (shown with SE). All DMSO vs. PladB-treated differences are significant (paired t-test; p<0.05) except when indicated (NS) **D.** γH2AX ChIP-qPCR data expressed as a ratio (PladB over DMSO) over a range of loci for asynchronous cells (top) and for cells synchronized in G1, S or G2/M (bottom). The DIvA cell line (yellow at right) in which site-specific breaks can be induced was used as a positive control for the experiment. The induction of γH2AX recruitment in DIvA cells was statistically significant (p<0.05). All other changes were not statistically significant.

We next asked if the loci experiencing γH2AX deposition at 24 hours would spatially overlap with DoG R-loops, as expected if such R-loops initiate DNA damage. For this, we performed chromatin immunoprecipitation (ChIP) experiments against γH2AX over multiple DoG R-loop hotspots. The DIvA/4-hydroxy tamoxifen (4OHT) system was used to induce double-strand breaks (DSBs) at predictable genomic loci (Iacovoni et al., 2010) as a positive control. ChIP-qPCR showed a 25-fold increased γH2AX enrichment in 4OHT-treated cells compared untreated cells (Figure 6D). By contrast, γH2AX was not increased over any locus four hours after PladB treatment. After 24 hours, while a trend towards increased γH2AX was observed, this trend was often not statistically significant and was not unique to DoG R-loop loci. Indeed, some RLL regions also showed a trend towards increased γH2AX (Figure 6D). Similar results were observed after cells were synchronized along the phases of the cell cycle. Taken together, these results suggest that, although PladB treatment induces DNA damage, such damage does not temporally or spatially correlate with regions that accumulate excessive R-loops.

### Depletion of the U2-associated Aquarius protein leads to readthrough transcription and DoG R-loops over stress-response genes

Aquarius (AQR) is a putative RNA helicase that physically associates with U2 snRNP components, shows distinctive binding to intronic branchpoints, and is required for efficient splicing *in vitro* and for tens of thousands of introns *in vivo* (De et al., 2015; Van Nostrand et al., 2020). Previous work showed that AQR depletion led to DNA damage and R-loop accumulation (Sollier et al., 2014). We therefore targeted AQR for siRNA depletion to determine if the PladB response was unique to that pharmacological agent or more generalizable to other U2 spliceosome-associated factors.

siRNA treatment led to a consistent 60% depletion in AQR levels after 72 hours which was accompanied by increased IR and by mild elevated DNA damage, as expected (Figure S7A-E). R-loop mapping using sDRIP-seq revealed that of the loci that exhibited significant changes, a clear majority of events (2:1) corresponded to R-loop losses (Figure 7A). Genes experiencing RLGs showed strong GO enrichment for ribosomal biogenesis and translation followed by RNA splicing (Figure 7B), which is similar to GOs identified for RLG or up-regulated genes upon PladB treatment (Figure 3D, F). RLG genes upon siAQR showed R-loop increases through gene bodies and downstream of genes (Figure 7C, D, Figure S7F) with more than half (53%) of RLG peaks mapped to DoG regions consistent with transcriptional readthrough. Only 10.5% of the thousands of retained introns observed upon AQR depletion intersected with RLGs, further suggesting that intron retention *per se* is not sufficient to drive R-loop formation. Instead, transcription perturbations such as DoG transcription and increased gene expression due to cellular stress responses drive the bulk of R-loop gains both upon PladB treatment and AQR depletion. AQR depletion also drove strong R-loop losses that were observed primarily through transcribed gene bodies (Figure 7E, Figure S7G) and generally consistent with a progressive loss of elongation.

**Figure 7.**
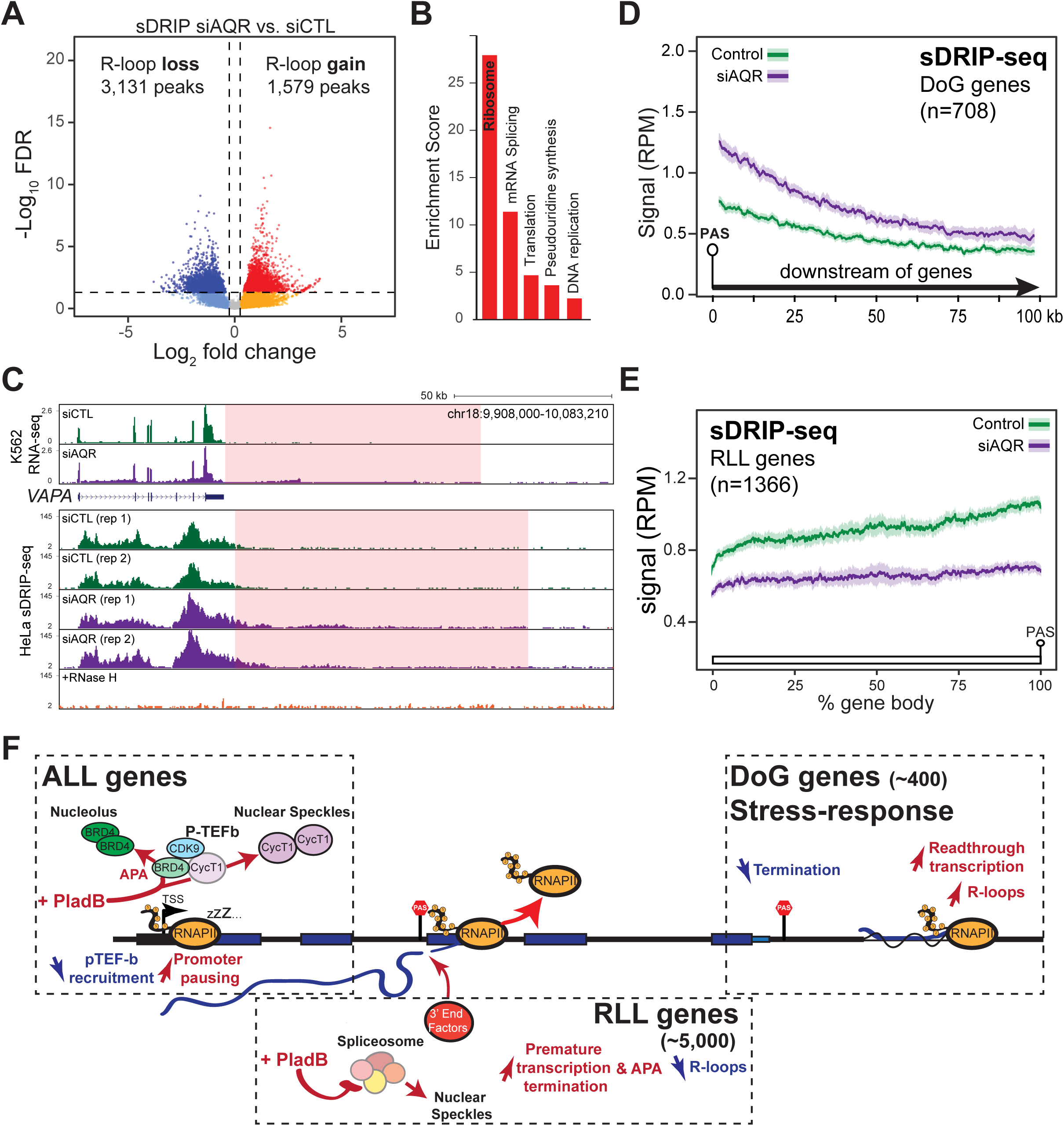
**A.** Volcano plots showing significant R-loop changes upon AQR depletion. **B.** Significant gene ontologies observed for RLG genes upon siAQR. **C.** Screenshot of RNA-seq (top) and R-loop distribution (bottom) for the *VAPA* gene upon AQR depletion. The highlighted region indicates significant gains in the region downstream of *VAPA*. **D.** Metaplot of R-loop signal downstream of the 708 genes found to gain R-loops upon AQR depletion. **E.** Metaplot of R-loop signal along the genes found to lose R-loops upon AQR depletion. **F.** Model. PladB treatment triggers reduction in pTEF-b recruitment and increase in promoter-proximal pausing for all genes, driven in part by the relocalization of cyclin T1 and BRD4. PladB treatment triggers increase in premature transcription termination and APA, resulting in a global decrease in R-loops at RLL genes. PladB also triggers decrease in termination, and readthrough transcription resulting in the formation of DoG R-loops for a subset of genes. See text for details.

## DISCUSSION

### SF3B1-mediated splicing inhibition triggers premature termination and promoter-proximal RNAP pausing

Pharmacological inhibition of the U2 spliceosome using PladB caused the retention of over 7,000 introns in mRNA pools (Figure 1), a likely underestimate of the impact of SF3B1 inhibition on splicing at the nascent RNA level (Drexler et al., 2020; Nojima et al., 2015; Sousa-Luis et al., 2021). In addition to these expected changes, PladB treatment caused a dramatic loss of transcription capacity affecting over 5,000 genes. We show that this results from the combined impact of at least two distinct mechanisms.

Increased promoter-proximal RNAP pausing (Figure 5), also observed independently (Caizzi et al., 2021), likely accounts for one such mechanism. Our observation that key pause-release factors, including BRD4 and cyclin T1, become sequestered into distinct nuclear sub-compartments away from transcriptionally active centers, provides a plausible explanation for this effect (Figure 7F). The relocalization of cyclin T1, a key component of pTEF-b, to splicing speckles is consistent with its less efficient recruitment to promoters (Caizzi et al., 2021). The relocalization of BRD4 to nucleoli is also expected to reduce its functional pools, further lowering pTEF-b recruitment and enhancing promoter-proximal pausing. Importantly, this effect does not appear specific to PladB as a similar accumulation of promoter-proximal transcripts was detected upon treatment of human HeLa cells with Isoginkgetin (IsoG) (Boswell et al., 2017), a non-SF3B splicing inhibitor that targets the U2 spliceosome at a later stage (O’Brien et al., 2008). Overall, increased promoter-proximal pausing due to U2 snRNP inhibition is likely to account for a significant reduction in gene expression given that the release of paused RNAP complexes is often a rate-limiting step (Adelman and Lis, 2012; Core and Adelman, 2019; Mayer et al., 2017). In addition, prolonged pausing can lower the release frequency and cause an overall reduction in RNA synthesis rates (Gressel et al., 2019; Gressel et al., 2017).

We show that, in addition, U2 snRNP inhibition activates global APA, leading to premature termination of these RNAP complexes that escaped promoter pausing. This second mechanism was evidenced by the steady decline of transcription levels through gene bodies and by the upregulation of thousands of annotated and novel alternative 3’-UTRs (Figure 4). Prior observations that PladB and other SF3B-targeted macrolides reduced the levels of elongation-associated Serine 2 phosphorylated RNAP and caused the dissociation of RNAP from chromatin (Koga et al., 2015) are consistent with our findings and suggest that these effects are again not specific to PladB. Indeed, treatment of human cells with IsoG, PladB, as well as other SF3B-targeted splicing inhibitors spliceostatin A and E7107, in addition to direct U2 snRNP depletion, were shown to cause reduced transcription (Boswell et al., 2017; Caizzi et al., 2021; Han et al., 2022) and induction of APA (Sousa-Luis et al., 2021). Interestingly, inhibition of the U1 snRNP complex also caused dramatic transcription elongation loss associated with premature termination (Berg et al., 2012; Kaida et al., 2010; Oh et al., 2017). Collectively, these results show that U1 and U2 snRNP inhibition trigger similar effects, confirming the tight integration of gene expression programs with the splicing machinery through diverse feedback mechanisms (Bentley, 2014; Herzel et al., 2017; Hsin and Manley, 2012).

While the effect of PladB on transcriptional dynamics appears to apply uniformly to all genes, the resulting impact on gene expression outputs varied depending on the intrinsic characteristics of the gene classes identified here (Figure 7). RLL genes suffered dramatic early elongation losses which dominated the PladB response landscape. The sensitivity of RLL genes to PladB can be traced to two distinguishing characteristics. First, RLL genes showed the lowest promoter-proximal RNAP store and the least increased pausing upon PladB treatment. Thus, any increased termination during elongation cannot be readily compensated by the release of paused RNAP complexes. Second, RLL genes carried cryptic PAS motifs shortly downstream of the TSS (median 3.7 kb) compared to other gene classes, ensuring early premature termination. In sharp contrast, DoG genes possessed large loads of paused RNAP even under control conditions, enabling them to maintain sufficient elongation capacity to complete their transcription. In addition, cryptic PAS sequences only occurred far downstream of the TSS (median 16.5 kb), ensuring that elongation was not interrupted prematurely. Unaffected genes, by virtue of their short size and strong gains of RNAP upon PladB treatment, also largely escaped elongation failure in response to PladB. Overall, the dramatic reduction of transcription capacity stemming from RLL genes accounts for the vast majority of co-transcriptional R-loop losses observed upon PladB treatment. This reinforces the notion that R-loop formation is a sensitive readout of transcription.

### PladB treatment causes readthrough transcription accompanied by *de novo* R-loop formation

PladB treatment also induced extensive readthrough transcription for a subset of 429 DoG genes (Figures 2, 3). Mechanistically, readthrough transcription may result from splicing alterations, consistent with the links between terminal exon splicing and termination (Davidson and West, 2013; Kyburz et al., 2006; Reimer et al., 2021; Rigo and Martinson, 2008, 2009). Readthrough may also result from a reduction of functional BRD4 pools given recent evidence that BRD4 coordinates the recruitment of 3’-RNA processing factors (Arnold et al., 2021). The characteristics of the DoG gene subclass are consistent with both possibilities. First, DoG genes show a propensity to undergo distal splicing alterations compared to unaffected genes, which may impact the mechanistic coupling between splicing and termination. In addition, DoG genes form only low levels of termination-associated R-loops compared to unaffected genes which may predispose them to lower termination efficiencies. Second, DoG genes, with their large stores of promoter-proximal RNAPs, are heavily regulated via pause release mechanisms and may therefore be most sensitive to a reduction in functional BRD4 levels. Importantly, readthrough transcription was also reported for HeLa cells treated with IsoG (Boswell et al., 2017) and upon AQR depletion (Figure 7), suggesting that it is not unique to PladB treatment and may represent a more universal response to U2 snRNP inhibition.

Almost all R-loop gain events upon PladB treatment mapped to DoG regions, delineating a novel class of aberrant *de novo* R-loops. Based on current understanding, DoG R-loops are likely to be similar in nature to other co-transcriptional R-loops, with the appearance of long DoG R-loops zones (Figure 2) due to the collective piling up of smaller sub-kilobase structures over R-loop prone regions (Malig et al., 2020). We note, however, that R-loop loads relative to nascent transcripts, appear significantly higher for DoG regions than for gene bodies (Figure S3D). This suggests that DoG transcription may be more prone to R-loop formation. Interestingly, DoG RNAs were previously shown to be nuclear-retained and likely chromatin-associated (Vilborg et al., 2015; Vilborg et al., 2017); it will be interesting to determine if R-loop formation underlies these characteristics. Collectively, excessive DoG R-loops occupied nearly 3 megabases of genomic space, although overall, R-loop losses overwhelmingly dominated over R-loop gains. These findings contrast with previous studies reporting that PladB and other splicing inhibitors including AQR depletion caused broad R-loops gains (Nguyen et al., 2018; Nguyen et al., 2017; Sollier et al., 2014; Wan et al., 2015). Such claims, which relied on S9.6-based imaging approaches, may be problematic given issues with S9.6 imaging (Crossley et al., 2021; Hartono et al., 2018; Phillips et al., 2013; Smolka et al., 2021). Overall, DoG R-loop gains occurred against the backdrop of a dramatic R-loop reduction genome-wide, itself driven by profound loss of transcription.

### PladB-induced “splicing shock” resembles other stress responses

DoG transcription has been observed in response to a range of cellular stresses including oxidative shock (Giannakakis et al., 2015; Vilborg et al., 2017), heat shock (Cardiello et al., 2018; Vilborg et al., 2017), osmotic shock (Rosa-Mercado et al., 2021; Vilborg et al., 2015; Vilborg et al., 2017), and viral infections (Bauer et al., 2018; Heinz et al., 2018; Hennig et al., 2018). When compared across stresses, DoG genes display both overlapping and specific gene ontologies that define the cellular response to a given stress (Morgan et al., 2022; Vilborg et al., 2017). DoG genes induced upon PladB treatment and AQR depletion were enriched for GOs related to protein translation, response to cellular stress, and mitochondrial energy homeostasis (Figure 3). Up-regulated genes identified by RNA-seq showed similar ontologies, suggesting that the cellular response to PladB is characterized by attempts at increasing translation capacity, mitochondrial respiration, and ATP synthesis in the face of dwindling and misspliced RNA transcript pools. A second emerging feature of several stress responses is that readthrough transcription occurs in the context of widespread transcriptional repression. Heat shock, for instance, caused decreased RNAPII occupancy across most protein-coding genes (Cardiello et al., 2018), accumulation of promoter-proximal paused RNAP, and broad gene down-regulation (Gressel et al., 2019; Mahat et al., 2016; Vihervaara et al., 2018). Similarly, osmotic stress led to reduced transcription for thousands of genes (Rosa-Mercado et al., 2021). As described above, PladB treatment also causes dramatic transcriptional shutdown due to increased promoter-proximal pausing and premature termination. Thus, SF3B-targeted splicing inhibition represents a novel example of a cellular stress that triggers induction of stress-responsive genes accompanied by readthrough transcription in the face of a global transcriptional shutdown.

One important aspect of the PladB stress response is the induction of alternative polyadenylation, leading to the increased usage of thousands of novel 3’-UTRs (Figure 5). This phenomenon may result from the functional coupling between splicing and 3’-RNA processing, whereby splicing loss unmasks cryptic PAS sites and provides access to the termination machinery (So et al., 2019). Inhibition of the PRMT1 arginine methyltransferase, which interacts with numerous RNA processing and splicing factors, also led to splicing disruptions and global APA induction (Giuliani et al., 2021). Similarly, evidence indicates that the transcriptional shutdown observed under heat shock involves increased premature termination (Cugusi et al., 2022). Interestingly, activation of APA is expected to lead to increased isoform switching both at the transcriptome and proteome levels. Whether such changes represent a maladaptive consequence of acute splicing inhibition or play a role in an adaptive stress response (Berkovits and Mayr, 2015; Mayr, 2019; Williamson et al., 2017) remains to be explored. Our observation that BRD4 and cyclin T1 are rapidly relocalized to nucleoli and nuclear speckles, respectively, indicates that protein trafficking may represent another dimension of stress responses. Heat stress indeed caused the rapid nucleolar localization of epigenetic regulators and BRD proteins (Azkanaz et al., 2019). The striking similarities between heat and osmotic stress responses to that induced by induced by targeting the splicing machinery using a pharmacological approach suggests that these environmental stressors may exert their effects on gene expression programs, at least in part, by impinging on the splicing machinery. In support of this, heat shock is known to trigger splicing inhibition from fly to mammalian cells (Shalgi et al., 2014; Yost and Lindquist, 1986) and SF3B1 itself was reported to be heat shock-sensitive (Kim Guisbert and Guisbert, 2017). Our work therefore defines a “splicing shock” stress response that is similar to several other cellular stress responses.

### Rethinking the links between splicing inhibition, genome instability, and aberrant R-loops

PladB treatment and AQR depletion trigger genome instability (Nguyen et al., 2017; Sollier et al., 2014; Wan et al., 2015). Mechanistically, it is commonly thought that excessive R-loops formed over retained introns underly these genome instability phenotypes. However, we show here that relatively few events of R-loop gains occur upon PladB treatment genome-wide and that less than 2 percent of retained introns overlapped with R-loop gains. Similar findings were observed for AQR depletion. We therefore conclude that inhibiting the U2 spliceosome does not trigger R-loop gains over retained introns. Given that PladB still allows binding of the spliceosome with its target pre-mRNAs (Effenberger et al., 2014), it is possible that RNA engagement is sufficient to prevent R-loop formation regardless of splicing activity, as proposed earlier (Bonnet et al., 2017). A similar logic may apply to AQR depletion given its putative role in defining the branchpoint in the context of the excised intron lariat, later in the splicing reaction process (Van Nostrand et al., 2020).

The identification of a novel class of DoG R-loops enabled us to test if these excessive R-loops were driving genome instability. Under our conditions, the DNA damage response was only detectable 12-24 hours post treatment, while DoG transcription was observed as early as 30 min (Figures 2, 6). The temporal disconnect between DoG R-loops and DNA damage markers was not due to cell cycle arrest or to DoG R-loops being cell cycle restricted; DoG R-loops were observed in G1, S, and G2 phases. Considering that toxic R-loops might require time to be processed into DNA breaks, we next tested if the γH2AX DNA damage marker, would at least show enrichment over regions that accumulate DoG R-loops. This was not the case. Instead, modest γH2AX gains were observed at some, but not all, DoG sites tested and over some RLL genes showing clear R-loop losses. The DNA damage response observed upon PladB treatment was therefore temporally and spatially uncoupled from DoG R-loop accumulation, raising questions as to whether excessive R-loops represent toxic genomic intermediates under splicing inhibition conditions.

Several non-exclusive alternative models can account for our observations. Transcriptional losses driven by PladB could for instance lead to progressive down-regulation of DDR genes (Figure 3), sensitizing cells to spontaneous genomic insults and ultimately resulting in DNA breaks. A similar mechanism was invoked in the context of the response to PRMT1 inhibition (Giuliani et al., 2021) and SF3B1 inhibition by E7107 (Han et al., 2022) and PladB (Pederiva et al., 2016). Conceptually, this predicts that genome instability events may be temporally and spatially uncoupled from R-loop events. Alternatively, it is possible that elongation-associated R-loops (Class II) measured by DRIP assays do not represent toxic structures and that another class of R-loops formed at paused promoters (Class I) drive the genomic conflicts from which DNA damage arises (Castillo-Guzman and Chedin, 2021). This is an attractive possibility given that Ribonuclease H1, which resolves R-loops *in vitro* and relieves DNA damage phenotypes when over-expressed, binds to paused promoters in a manner that is dynamically coupled to pausing (Chen et al., 2018; Chen et al., 2017). Our observation that PladB treatment causes increased promoter pausing suggests that promoter-bound Class I R-loops may also increase and lead to DNA damage. Increased Class I R-loops were shown to lead to increase promoter DSBs upon depletion of the R-loop unwinding DDX41 helicase (Mosler et al., 2021). Overall, our findings put into question the model that proper splicing is necessary to protect the genome from the damaging consequences of excessive genic R-loops caused by retained introns (Li and Manley, 2005; Paulsen et al., 2009; Tresini et al., 2015). Our study also highlights the necessity of directly characterizing R-loop patterns at the genome scale and of factoring the dynamic cellular and transcriptional responses that likely accompany any genetic or pharmacological perturbation targeted towards RNA processing factors.

## Limitations of the study

This study was primarily focused on the response to the PladB inhibitor, which targets the U2 spliceosome through the SF3B complex, and to AQR depletion. It will be important to establish if similar effects would result from targeting the variety of additional splicing-related factors previously associated DNA damage phenotypes upon their depletion (Paulsen et al., 2009). In addition, we did not directly assess the distribution of DNA breaks that result from PladB treatment, thereby limiting our ability to conclude about the mechanisms by which splicing dysfunction ultimately result in genome instability. Addressing this point will require precise mapping of DNA breaks.

## Acknowledgements

Work in the Chedin lab is supported by the National Institutes of Health grand R35 GM139549 (to F.C). D.C.G. was supported in part by the NIH T32 predoctoctoral training program in Molecular and Cellular Biology (GM007377) and by an NIH F31 individual fellowship (GM136143). The DIvA cell line was a kind gift from Dr. Giovanni Tonon (San Raffaele Scientific Institute, Milan, Italy). Hydra RNA was a kind gift from Dr. Celina Juliano. We thank members of the Chedin lab for useful suggestions.

## Authors contributions

D.C.G performed all PladB experiments with the assistance of L.A.S for R-loop mapping and T.S. for immunofluorescence microscopy. S.R.H. performed computational data analysis with help from D.C.G. M.F. performed APA validation experiments and all experiments related to AQR. F.C. conceived and supervised the project, analyzed data, and wrote the manuscript along with all authors.

## Data availability

All high-throughput genome sequencing datasets are available on the NCBI GEO website under accession number GSE148768.

## STAR* Methods

### Analysis of transcriptome alterations by RNA-seq and RT-qPCR

Human K562 cells were treated with PladB (100 nM final concentration) for 2 and 4 hours, at which point they were harvested. Mock-treated (DMSO) and untreated samples were also collected as controls. At each time point, total RNA was isolated from 1x10^6^ cells using TRI reagent (Life Technologies, Grand Island, NY) and Direct-zol RNA MiniPrep kit (Zymo Research). Isolated RNA was then treated with 2.5 µl of DNase I (NEB) for 1 hour at 37°C. Following DNase I treatment, RNA was cleaned up using RNA Clean & Concentrator-5 Kit (Zymo Research) and resuspended in 15 µl. After cleanup, poly(A) RNA was selected and RNA-seq libraries were constructed using the Illumina TruSeq kit prior to sequencing on an Illumina HiSeq 4000 instrument. Mapping and all downstream analysis were performed on two independent biological replicates.

For RT-qPCR, cDNA was generated using iScript Reverse Transcription Supermix for RT-qPCR (Bio-Rad), with 900 ng of human RNA and 100 ng of Hydra RNA (as exogenous normalizer). cDNA was diluted 1:3 (final volume 60 µl). For RT-qPCR, 1 µl of 1:3 dilution of the cDNA was used per well in a 20 µl reaction, along with 2 µl of 10 µM primer sets, and 10 µl of SsoAdvanced Universal SYBR Green Supermix (Bio-Rad). Reactions were run in triplicate on a CFX96 Touch Real-Time PCR Detection System (Bio-Rad) with the following protocol: 95°C (30 sec), 40 cycles of 95°C (10 sec) and 60°C (30 sec), followed by melt curve analysis (65-95°C). Quantification was calculated using the ΔΔC_t_ method. All values were normalized to a Hydra gene (RP49) and to time zero (Untreated/DMSO). RT-qPCR was performed on three independent biological replicates.

### Analysis of R-loop genomic distribution by DRIP-seq and DRIP-qPCR

PladB treated and control K562 cells were harvested 2 and 4 hours after PladB treatment. At each time point, genomic DNA was isolated from 5x10^6^ cells and extracted as described (Sanz and Chedin, 2019). S9.6-based DNA:RNA hybrid immunoprecipitation (DRIP) was performed as described previously (Ginno et al., 2012). In brief, 4.4 μg of restriction digested genomic DNA was immunoprecipitated for 16 hours at 4°C with 10 μL of S9.6 antibody. Three immunoprecipitated samples were combined into one for library construction. Once the quality of the combined immunoprecipitation was checked by qPCR at a range of positive and negative test loci, barcoded sequencing libraries were built, library quality was verified, and sequencing was performed on Illumina HiSeq or NovaSeq instruments. Mapping and all downstream analysis was performed on two independent biological replicates. R-loops were also mapped using sDRIP-seq, a variant of DRIP-seq that permits strand-specific R-loop mapping after genome fragmentation using sonication (Smolka et al., 2021).

For DRIP-qPCR, 2 μl of immunoprecipitated DNA (and 1:10 diluted input DNA) was used per well in a 20 μl reaction, along with 2 µl of 10 µM primer sets, and 10 µl of SsoAdvanced Universal SYBR Green Supermix (Bio-Rad). Reactions were run in duplicate on a CFX96 Touch Real-Time PCR Detection System (Bio-Rad) with the following protocol: 95°C (30 sec), 40 cycles of 95°C (10 sec) and 95°C (30 sec), followed by melt curve analysis (65-95°C). All values were normalized relative to input and normalized to Control (Untreated & DMSO) to calculate fold enrichment. DRIP-qPCR was performed on three independent biological replicates. For RNase H treatment, 6 µg of restriction digested DNA was treated with RNase H1 (Loomis et al., 2014) for 1 hour at 37°C using a 1/100 dilution of the stock enzymes in 50 mM Tris-HCl pH 7.5, 75 mM KCl and 3 mM MgCl2. DNA was then purified with phenol/chloroform and ethanol precipitated before being used in S9.6 immunoprecipitations as described above.

### Analysis of nascent transcripts using EU-seq

Human K562 cells treated or not with PladB were incubated with 5-ethynyl uridine (EU) (0.5 mM final concentration) for 20 min prior to harvesting two- and four-hours post PladB treatment. At each time point total RNA was isolated from 3x10^6^ cells and initially processed as described for RNA-seq. EU-RNA was enriched using Click-iT Nascent RNA Capture Kit (Invitrogen), with 5 µg of EU-RNA for biotinylation by click reaction and 1 µg of biotinylated RNA for binding to Streptavidin T1 magnetic beads. cDNA was generated using iScript Select cDNA Synthesis Kit (Bio-Rad). cDNA was cleaned up before second-strand synthesis. Then cDNA was ligated to Illumina barcoded adapters for sequencing. Library quality was checked on an Agilent BioAnalyzer and sequencing were performed on Illumina HiSeq or NovaSeq instruments. Mapping and downstream analysis was performed on two independent biological replicates.

### Sequencing data processing and computational data analysis

STAR version 2.7 (2.7.0f_0328) was used to map all sequencing reads and splice junctions (default parameters). Macs2 v2.2.5 was used to call EU-seq and sDRIP-seq signals according to defined bins. For DRIP-seq, we used restriction digest as bins. For RNA-seq, we used annotated genes and gene features (e.g. exon, intron) as their bins using Gencode version 19 (GRCh37). For subsequent analysis, the gene set was further filtered for highest APPRIS principal protein-coding genes with expression levels of at least 10 RPKM, as an annotation file. Htseq-count v0.11.2 was then used to count reads into their bins. DESeq2 v1.24 was used to calculate significant gains or losses (adjusted p-value of ≤ 0.05 and |log2(FC)| ≥ 1). rMATS v4.0.2 were used to identify significant changes (adjusted p-value of ≤ 0.05 and |log2(FC)| ≥ 0.32) in skipped exons (SE), A3SS, A5SS, and intron retentions (IR). In addition to rMATS, intron retention events were called by identifying significant increases in intronic reads using DESeq2 as well as splice junctions from STAR. To distinguish IR events from other possible events, intronic gains that spanned less than 75% of intron length were categorized into putative 3’UTRs if their 5’ end were adjoined (within +/- 10% of intron length) to the 3’ end of an exon or junction, and their 3’ end were not adjoined to the 5’ end of an exon or junction (and vice versa for putative 5’UTRs). For location analysis, we divided the genic space into several regions: promoters (+/-1kb of TSS), gene body (+1kb of TSS to -1kb of PAS), terminal (+/- 1kb of PAS), downstream of genes (DoG), and intergenic regions (everything else). DoG regions were defined as regions between a gene’s terminal region (+1kb of PAS + 1 kb) up to either 150 kb downstream if no neighbor was found prior, or to the promoter of its closest downstream gene minus 1 kb, whichever is closest. The 150 kb limit was chosen because a few test genes showed DoG regions extending more than 100 kb. Post-analysis, very few genes had DoG regions longer than 100 kb. To correctly assign a DoG region to the gene from which it originated, we annotated the strandedness of EU-seq and DRIP-seq signals based on the ratio of positive and negative strands of RNA-seq and sDRIP-seq signals.

### Fixation and labelling for γH2AX, BRD4, SF3B2 and AQR immunofluorescence

HeLa cells were grown, fixed, permeabilized, washed, immunostained, and imaged in 35 mm glass bottom poly-D-lysine-coated dishes (P35GC-1.5-14-C, MatTek) using 2 mL volumes of media and buffer solutions. All steps for fixation and immunofluorescence were carried out at room temperature. Cells were fixed in freshly prepared 1% formaldehyde in PBS for 10 minutes. Fixation solutions were quenched with the addition of 200 µL of 1 M glycine in PBS. Samples were washed once with PBS, and then incubated in permeabilization buffer (PBS with 0.1% Triton X-100) for 10 minutes. Samples were then incubated in staining buffer (TBST with 0.1% BSA (A9647-50G, Sigma)) for 10 minutes with rocking and a 1:1000 dilution of primary anti-phospho-Histone H2A.X (Ser139) antibody (05-636, Sigma) or anti-IBP160 (AQR) (A302547A, Bethyl Laboratories) was added for 1 hour with rocking. For SF3B2 and cyclin T1 or BRD4 samples: a 1:500 dilution of primary anti-SAP-145 antibody (sc-513930, Santa Cruz) and 1:1000 dilution of primary anti-BRD4 antibody (702448, Invitrogen) or 1:1000 dilution of primary anti-cyclin T1 antibody (PA5-77892) was added for overnight incubation with rocking. Samples were then washed once with staining buffer and incubated with a 1:2000 dilution of secondary anti-mouse AlexaFluor 488 conjugate (A28175, Invitrogen) and/or AlexaFluor 594 (A32740, Invitrogen) for 1 hour with rocking with samples kept concealed from light from this step onward. A 2.5 µg/mL DAPI solution in staining buffer was then added for 2 minutes and washed in TBST for 10 minutes with rocking. Samples were then immediately imaged. For each experiment, all samples were prepared, treated, and stained in parallel from one master antibody and/or dye dilution to ensure uniform treatment and staining efficiencies. Fixation and labelling was performed for two independent biological replicates.

### Imaging and image analysis

Fixed cell imaging was performed using a confocal laser scanning biological microscope [Olympus FV1000] with a 60X objective. For each experiment, all conditions were imaged in parallel with identical exposure times and laser settings. Images were analyzed and quantified using ImageJ. Statistics and data visualization were done using RStudio. P-values were determined by a Wilcoxon Mann-Whitney test using wilcox.test() function. The number of foci were counted for individual nuclear regions defined by thresholding the DAPI channel to obtain Regions of Interest (ROI) for analysis. ROIs containing more than one nucleus were removed from downstream analysis. Find Maxima was applied to the 488 channel using a noise tolerance of 700 to generate a binary image of local maxima (γH2AX foci). Analysis was then performed on nuclear ROIs to quantify the number of γH2AX foci per nucleus. For BRD4, SF3B2 and AQR IF analysis, total nuclear intensity was measured for each ROI.

### Analysis of γH2AX genomic distribution using ChIP-qPCR

Human K562 cells (1x10^7^ cells) were treated with PladB (100 nM final concentration) for 4 and 24 hours or DMSO as mock-control. DIvA cells were treated with or without 4-hydroxy tamoxifen (4OHT) (300 nM final concentration) for 4 hours. K562 cells were then collected and resuspended in PBS, then formaldehyde was added to a final concentration of 1% and crosslinking was allowed to proceed for 10 minutes at room temperature, while the samples were rocking. For DIvA cells, media was removed and replaced with PBS and Formaldehyde was added to dish. In order to stop the reaction, fresh glycine was added to a final concentration of 0.125 M. For DIvA cells, the cells were then scraped and washed once with PBS. After 10 minutes cells were washed once with PBS. Pelleted cells were then resuspended in RIPA lysis buffer (150 mM NaCl, 5 mM EDTA, 50 mM Tris pH 8.0, 1% IGEPAL CA-630, 0.5% sodium deoxycholate, 0.1% SDS) supplemented with phosphatase and protease inhibitors (PHOSS-RO, Sigma and 539134, Millipore). The lysate was then sonicated 30 secs for 15 cycles (Diagenode Bioruptor) to obtain DNA fragments of about 500-1000 bp. Samples were then centrifuged for 15 mins at 4°C at 15,000 rpm and the supernatants were transferred to new tubes. Samples were then diluted in RIPA buffer (100 µl chromatin : 900 µl RIPA buffer) and phosphatase and protease inhibitors were added to samples. A portion of the sample was removed and saved as Input. The rest of the sample was subjected to a 60 min preclearing with 60 µl of previously blocked protein A/G magnetic agarose beads (PI78609, Thermo Scientific) at 4°C. Blocking was achieved by incubating the agarose beads in RIPA buffer with 0.1% BSA for 30 minutes. Precleared samples were incubated with rotation overnight at 4°C with either 10 µg of anti-phospho-Histone H2A.X (Ser139) antibody (05-636, Sigma) or 10 µg of IgG as control. Immune complexes were then recovered by incubating the samples with 60 µl of previously blocked protein A/G magnetic agarose beads for 1 hour at 4°C on a rotating wheel. Beads were washed 6 times in wash buffer (150 mM NaCl, 10 mM Tris pH 7.5, 2 mM EDTA, 0.1% IGEPAL CA-630) for 10 minutes followed by an additional wash with 10 mM Tris-EDTA pH 8.0. Elution and crosslink reversal were achieved by incubating samples in 10 mM Tris-EDTA + 0.1% SDS and proteinase K at 65°C for 4 hours. Elutions were then transferred to a phase-lock separation tube. The beads were then washed one more time with 10 mM Tris-EDTA and 0.5 M NaCl for 10 minutes at 65°C and the supernatant was added to the phase-lock separation tube. DNA was extracted by the phenol-chloroform method and precipitated with ethanol.

For ChIP-qPCR, 2 μl of immunoprecipitated DNA (and 1:4.5 diluted input DNA) was used per well in a 20 μl reaction, along with 2 µl of 10 µM primer sets, and 10 µl of SsoAdvanced Universal SYBR Green Supermix (Bio-Rad). Reactions were run in duplicate on a CFX96 Touch Real-Time PCR Detection System (Bio-Rad) with the following protocol: 95°C (30 sec), 40 cycles of 95°C (10 sec) and 95°C (30 sec), followed by melt curve analysis (65-95°C). All values were normalized relative to input. ChIP-qPCR was performed on two independent biological replicates for K562 samples and two technical replicates for DIvA samples.

### Cell cycle synchronization

To obtain G1 stage-specific synchronization, sub confluent K562 cells were treated for 24 hours with thymidine (2 mM) and released into pre-warmed medium for 3 hours before adding nocodazole (100 ng/mL) for 12 hours. Then the cells were released for either 8 or 10 hrs before adding PladB (100 nM) for 2 or 4 hours, respectively. The cells were then collected for both flow cytometry and DRIP-qPCR. Each experiment was performed 2 times independently. To obtain S stage-specific synchronization, sub confluent K562 cells were treated for 24 hours with thymidine (2 mM) and released into pre-warmed medium containing PladB (100 nM) for 2 or 4 hours. The cells were then collected for both flow cytometry and DRIP-qPCR. Each experiment was performed 2 times independently. To obtain G2/M stage-specific synchronization, sub confluent K562 cells were treated for 24 hours with thymidine (2 mM) and released into pre-warmed medium for 6 hours before adding RO-3306 (10 mM) and PladB (100 nM) for 2 and 4 hours. The cells were then collected for both flow cytometry and DRIP-qPCR. Each experiment was performed 2 times.

### Determination of Cell Cycle Profiles

2x10^6^ K562 cells were washed twice and resuspended in 1 mL of PBS. This mixture was slowly added to 4 mL of 100% ethanol and cells were incubated on ice for 30 minutes. The fixed cells were centrifuged at 800xg for 5 minutes and washed once with PBS before staining with 600 ul of Propidium Iodide (PI). The samples were detected with an BD FACS Canto II Flow Cytometer. Cell-cycle distribution was analyzed using the FlowJo Software with a minimum of 10,000 cells. Each experiment was performed 2 times independently.

### siRNA depletion of AQR

HeLa cells were plated at 25,000 cells/ml in antibiotic free DMEM 24 hours prior to transfection. Cells were transfected with either a 20 nM final concentration of siAQR (s18725, Thermo Fisher) or siControl (4390843, Ambion) using DharmaFECT (Horizon). siRNA was diluted in 50ul of Opti-MEM (Thermo Fisher) as was DharmaFECT and incubated for 5 minutes at room temperature. The two mixtures were combined by gently pipetting then incubated at room temperature for 20 minutes. The DhamaFECT and siRNA mixture was added dropwise to the cells. Cells were harvested for sDRIP, immunofluorescence and RT-qPCR 72 hours after transfection. All assays were performed as described above.

---

**Figure S1:**
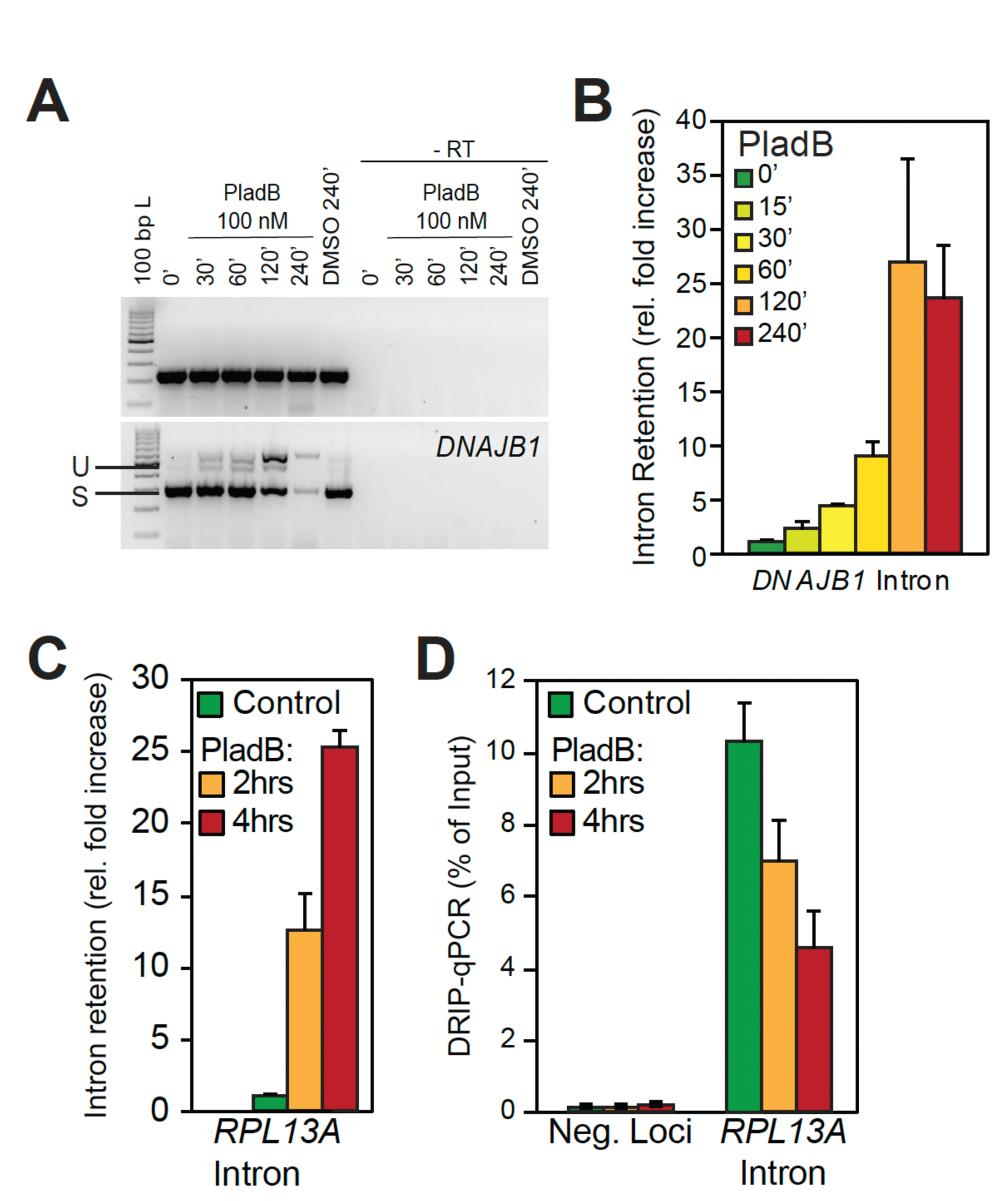
**A.** Gel-based RT-PCR splicing assay verifies that PladB treatment causes intron retention over the *DNAJB1* target gene, but not the control *ACTB* gene. No PCR products were obtained if reverse-transcription (RT) was omitted, as expected. **B.** RT-qPCR assay confirms that PladB treatment causes intron retention for *DNAJB1* target gene. IR starts to occur as early as 15 min. **C.** RT-qPCR assay confirms that PladB treatment causes intron retention for *RPL13A*, a positive R-loop forming gene. **D.** Bar chart of DRIP-qPCR (as percent input) for *RPL13A*. Each bar is the average of three-independent experiments (shown with SE). PladB treatment results in loss of R-loops over intron with IR.

**Figure S2:**
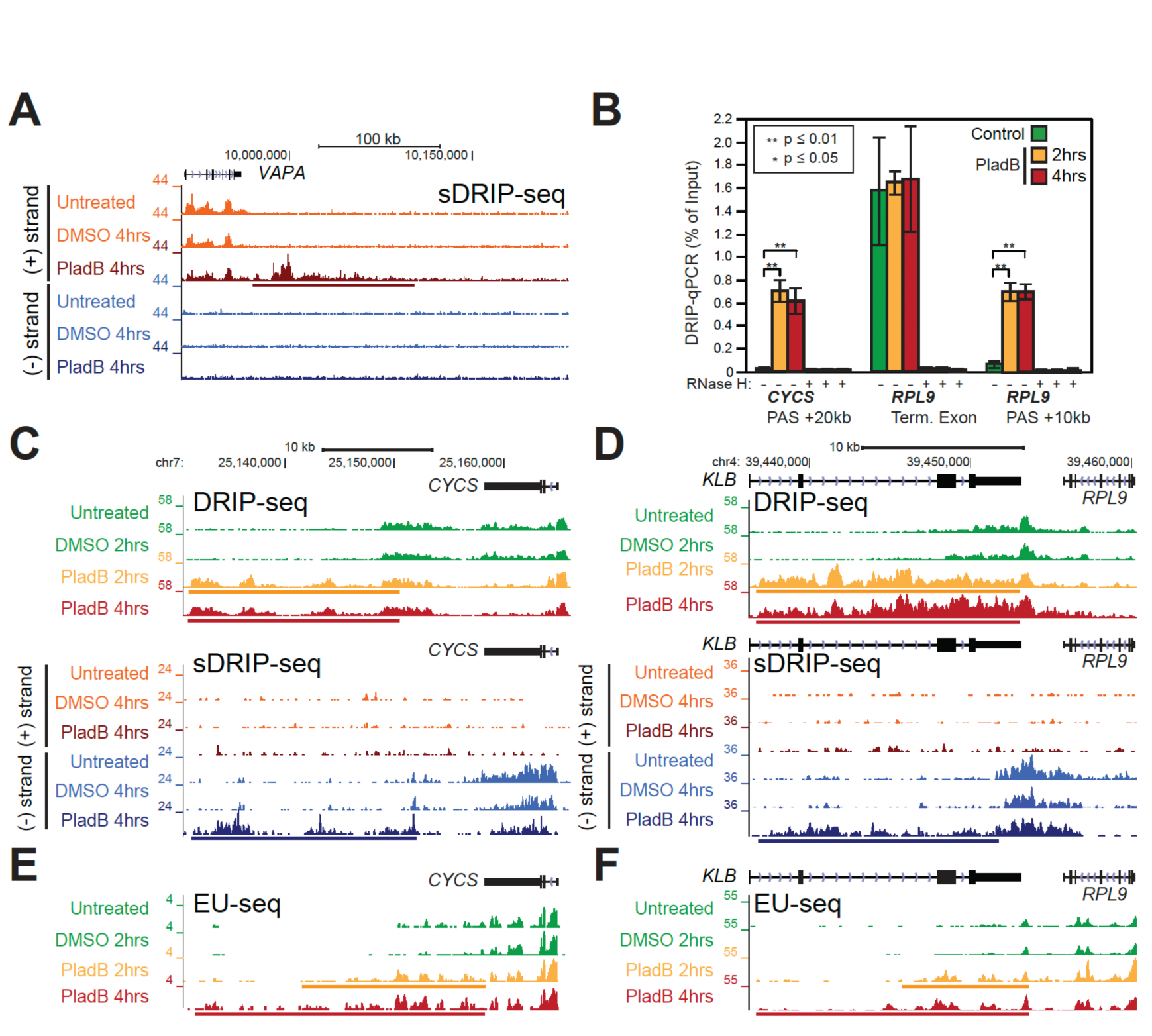
**A.** Genome browser screenshot over the *VAPA* gene and ∼200 kb downstream showing plus and minus strand sDRIP-seq signal obtained from controls and PladB-treated K562 samples. R-loop gains occur directly downstream of gene (DoG) only on the plus strand (colored bar). **B.** Bar chart of DRIP-qPCR (as percent of input) for terminal and DoG regions. Each bar is the average of three-independent experiments (shown with SE). **C.** Genome browser screenshot over the *CYCS* gene and downstream region showing DRIP-seq (Top) and sDRIP-seq signals over the plus and minus strands (Bottom) from both controls and PladB-treated K562 samples. R-loop gains occur directly downstream of gene (DoG) and only on the minus strand (colored bar). **D.** Genome browser screenshot over the *RPL9* gene and downstream region showing DRIP-seq (Top) and sDRIP-seq signal over the plus and minus strands (Bottom) from controls and PladB-treated K562 samples. R-loop gains occur directly downstream of gene (DoG) and only on the minus strand (colored bar). **E.** Genome browser screenshot showing the same region as in (B) but now displaying EU-seq signals. Region with increased R-loops also have increased nascent transcripts. **F.** Genome browser screenshot showing the same region as in C but now displaying EU-seq signals. Region with increased R-loops also have increased nascent transcripts.

**Figure S3:**
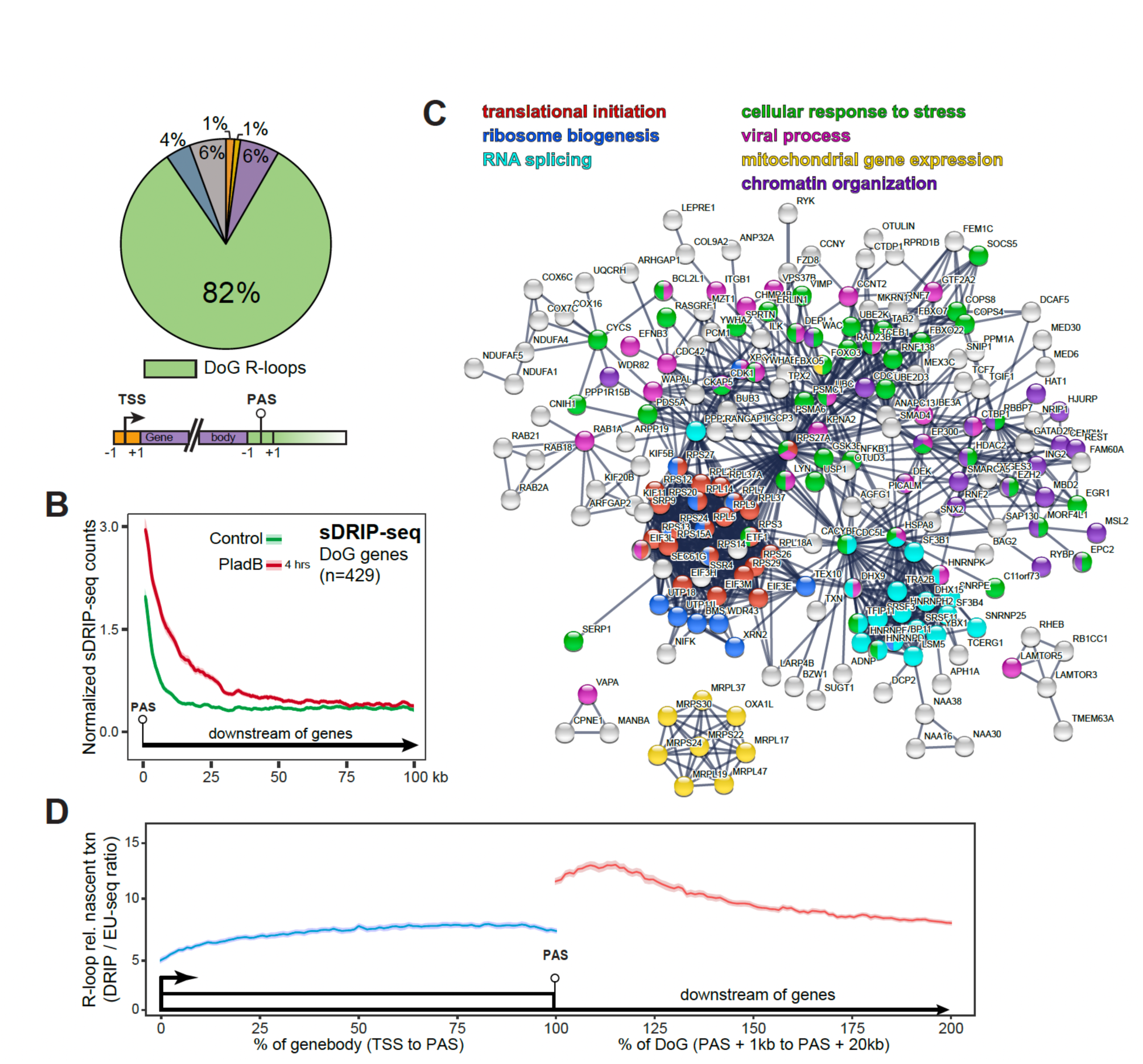
**A.** Location analysis of R-loop peak gains for PladB-treated K562 sample (4 hours) according to downstream of gene (DoG) annotation pipeline. Annotation scheme is below. 82% of R-loop gains 4 hours post-PladB treatment specifically map to downstream of gene (DoG) regions. **B.** Metaplots of sDRIP-seq signals extending from nearest PAS for all annotated DoG genes. For each sample, the signal is shown as a trimmed mean (line) surrounded by SE (shaded). **C.** STRING interaction analysis. All 429 DoG genes were used as input for STRING analysis and a network was built based on highest confidence (0.9). Nodes are color coded based on biological processes. Nodes that had no interaction are not shown. **D.** Scaled metaplot of R-loop signal (DRIP-seq) relative to transcription levels (EU-seq) is graphed for gene body and DoG regions for DoG genes. Data is shown as the trimmed mean (line) along the standard error (shaded).

**Figure S4:**
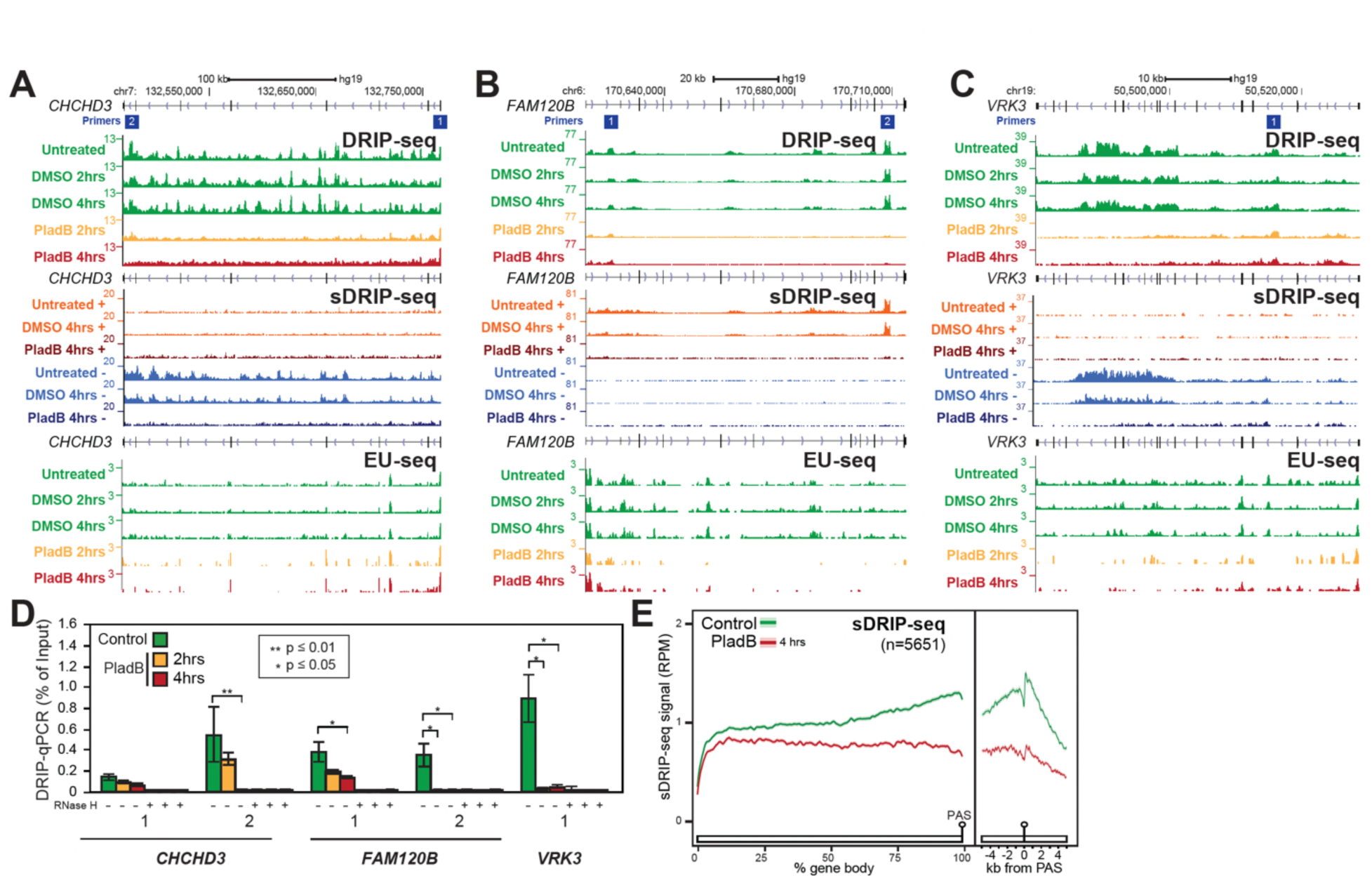
A-C. Genome browser screenshots over the *CHCHD3, FAM120B,* and *VRK3* genes showing DRIP-seq signal (Top), sDRIP-signal (Middle), and EU-seq signal (Bottom) obtained from controls and PladB-treated K562 samples. R-loop loss is directional with transcription. **D.** Bar charts of DRIP-qPCR (as percent of input) for *CHCHD3, FAM120B,* and *VRK3* validating R-loop losses. Each bar is the average of three-independent experiments (shown with SE). Primer locations are indicated on each panel. P-values from paired t-tests are indicated. **E.** Metaplot of sDRIP-seq signals over gene body (as % of gene length) and terminal regions (+/- 5kb from PAS) of RLL genes. Control and PladB-treated samples are color-coded as indicated. For each condition, the signal is shown as a trimmed mean (line) surrounded by the standard error (shaded).

**Figure S5:**
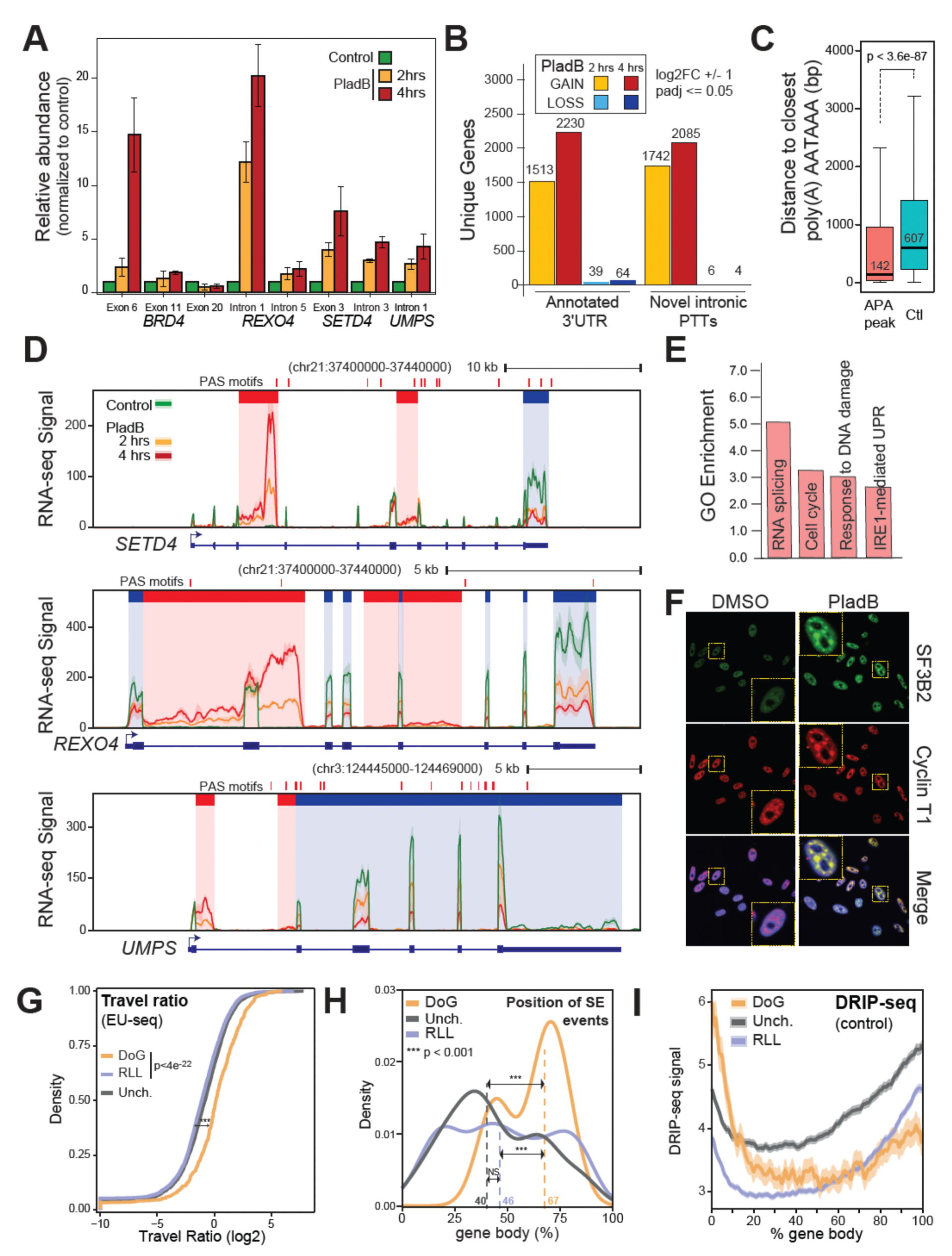
**A.** Bar chart of RT-qPCR (as fold change over control) at indicated PladB-induced APA sites loci after PladB treatment. Each bar represents the average of three independent experiments (shown with SE). **B.** Systematic annotations of putative APA sites 4 hours after PladB treatment. Events were broken down between arising over previously annotated 3’UTRs and novel intronic 3’-UTRs. **C.** Distance between the end of novel intronic APA regions and annotated polyA motifs. Control distances were calculated after shu_ling the position of the APA regions within the same genes. P-value was calculated using a Wilcoxon test. **D.** RNA-seq signal for *REXO4*, *SETD4*, and *UMPS* under control and PladB-treated conditions. Highlighted in red and blue are sites of significantly increased or decreased APA usage. **E.** Enriched gene ontologies for Plad-induced APA genes. **F.** Representative immunofluorescence microscopy images from DMSO- and PladB-treated (4 hours) HeLa cells analyzed for SF3B2 and Cyclin T1 distribution and DAPI. Cyclin T1 becomes relocalized to splicing speckles. **G.** Travel ratios for DoG, RLL and una_ected genes are plotted using EU-seq signals under control conditions. Data is shown as the trimmed mean (line) along the standard error (shaded). **H.** Distribution of skipped exon events along DoG, una_ected, and RLL genes is plotted along gene bodies 4 hours post PladB treatment. The dashed lines indicate the position of the median position of such events. All p-values were determined by a Wilcoxon Mann-Whitney test (one, two, and three stars indicate p-value of ≤ 0.05, 0.01, and 0.001, respectively; NS: Not Significant). **I.** Metaplot of DRIP-seq signals along DoG, una_ected, and RLL genes were plotted along gene bodies under control conditions. Data is shown as the trimmed mean (line) along the standard error (shaded).

**FIGURE S6:**
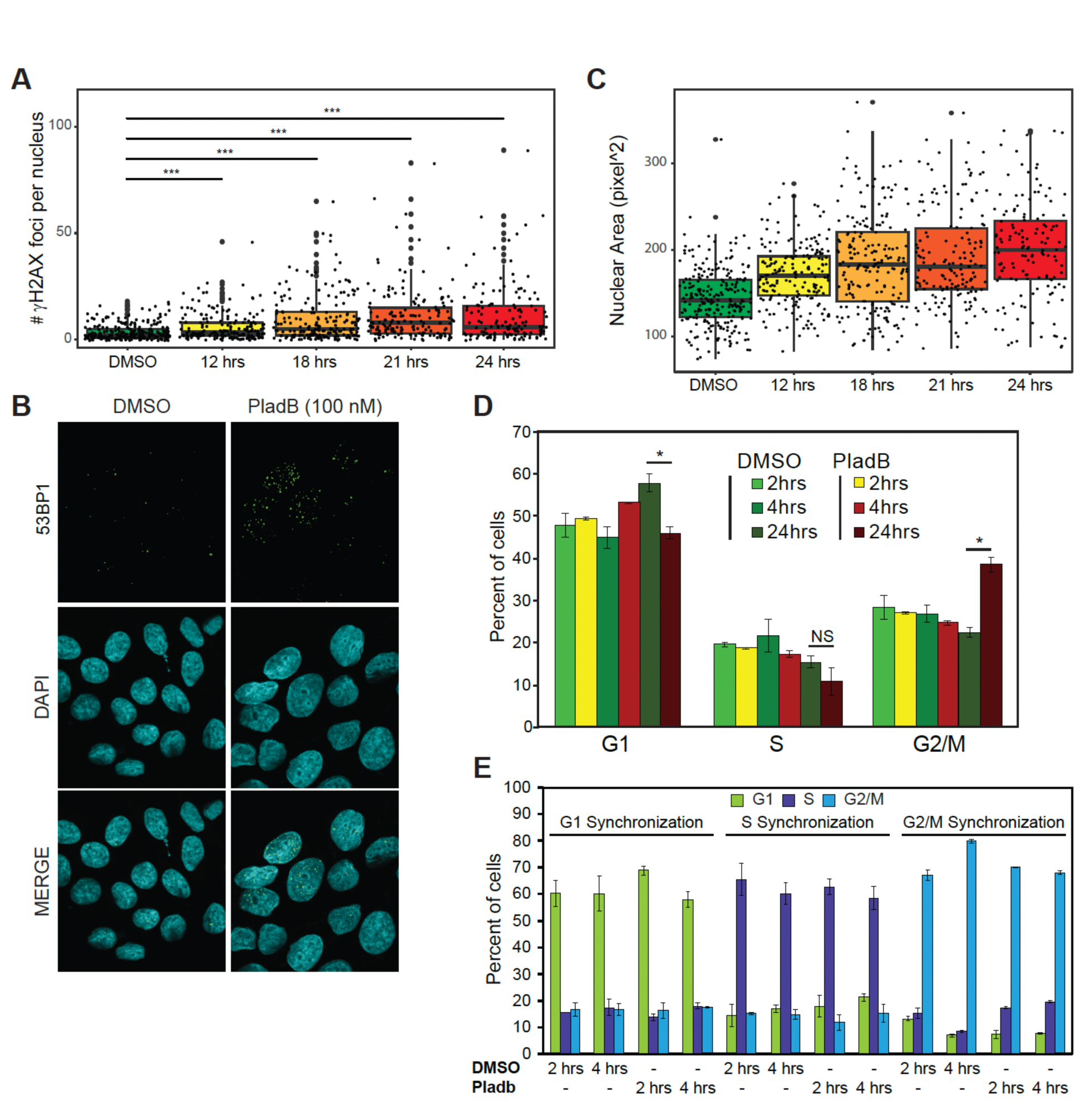
**A.** Box plot of γH2AX foci over time in HeLa cells treated with 100 nM PladB or DMSO at the indicated timepoints. *** indicates p-value < 0.001. Each dot corresponds to one cell collected over two independent biological replicates. **B.** Representative immunofluorescence microscopy images from DMSO- and PladB-treated (24 hours) HeLa cells analyzed for the 53BP1 DNA damage marker and DAPI. **C.** Box depicting the nuclear area of HeLa cels treated with 100 nM PladB at the indicated timepoints. **D.** Cell cycle distribution by PI staining: Bar plot representing the percentage of cells in G1, S, and G2/M in K562 cells at indicated times after DMSO or PladB treatment. Bars represent an average of 4 independent measurements, Error bars represent SE. **E.** FACS analysis of cell cycle distribution post synchronization verifying that cells were properly synchronized.

**FIGURE S7:**
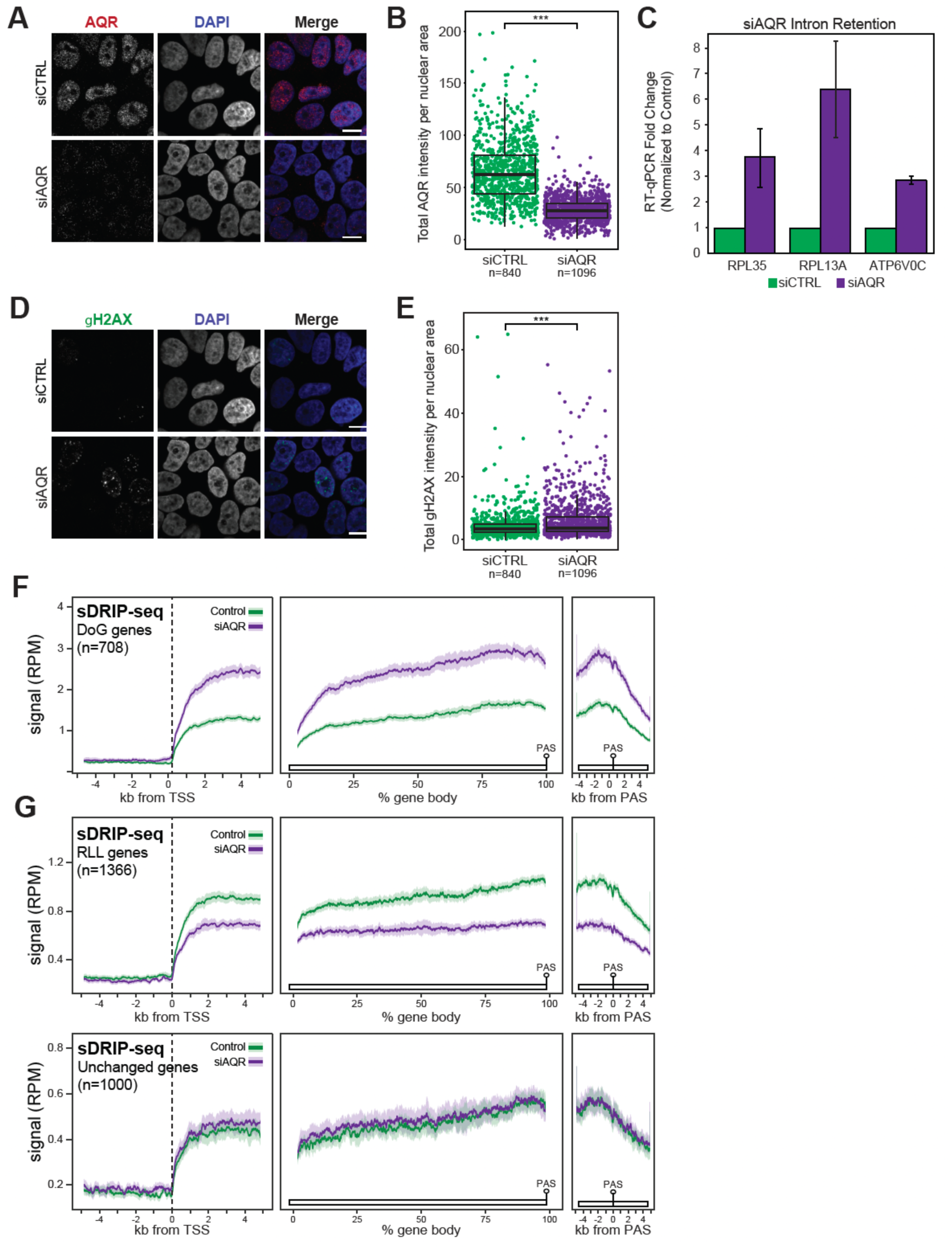
**A**. Immunofluorescence imaging of HeLa cells treated with mock- or AQR-targeted siRNAs. **B**. Quantification of AQR signal over the indicated number of cells showing significant AQR depletion. **C**. Verification of intron retention using RT-qPCR upon siAQR treatment at three distinct genes. **D**. Immunofluorescence imaging DNA damage marker γH2AX in HeLa cells treated with mock- or AQR-targeted siRNAs. **E**. Quantification of γH2AX signal over the indicated number of cells showing significant DNA damage induction. **F-G**. Metaplots of R-loop signal over RLG genes (F), RLL genes (G – top) and unchanged genes (G – bottom).

